# Cytoplasmic incompatibility between Old and New World populations of a tramp ant

**DOI:** 10.1101/2020.03.13.988675

**Authors:** Çiğdem Ün, Eva Schultner, Alejandro Manzano-Marín, Laura V. Flórez, Bernhard Seifert, Jürgen Heinze, Jan Oettler

## Abstract

As we enter the Anthropocene, the evolutionary dynamics of species will change drastically, and as yet unpredictably, due to human activity. Already today, increases in global human traffic have resulted in the rapid spread of species to new areas, leading to the formation of geographically isolated populations. These go on to evolve in allopatry, which can lead to reproductive isolation, and potentially, the formation of new species. Surprisingly, little is known about such eco-evolutionary processes in ants, even though numerous invasive ant species are globally distributed in geographically isolated populations. Here, we describe the first case of cytoplasmic incompatibility (CI) between populations of a cosmotropic distributed tramp ant with Asian roots, *Cardiocondyla obscurior,* which has acquired a novel *Wolbachia* strain in the New World. Our study uncovers the first symbiont-induced mechanism of reproductive isolation in ants, providing a novel perspective on the biology of globally distributed ants.

## Introduction

Ants are the most abundant group of insects on earth, and numerous ant species are classified as highly invasive on a global scale. Their distribution is, on the one hand, strongly facilitated by humans. This is evidenced, for instance, by the rapid worldwide spread of the Argentine ant and the fire ant from their origin in South America^1,2^. On the other hand, the particular biology of ants, characterized by reproductive division of labor between highly fecund queens and sterile workers, allows the establishment of large populations from just one founding propagule. Introduced populations thus go through a genetic bottleneck, reducing diversity within the population while at the same time increasing diversity between populations^3^. Such populations then evolve in allopatry, and differences caused by genetic drift may be further amplified over time by selection, eventually leading to reproductive isolation. Reproductive isolation, in turn, together with the evolution of mating systems, underlies the formation of new species and, hence, the vast diversity of life. By allowing ants (and other organisms) to establish allopatric populations around the world, humans have inadvertently created a valuable scenario for studying species evolution in real time.

Reproductive isolation between populations that live in allopatry can evolve in the form of post-mating mechanisms of reproductive isolation (PoMRI). Importantly, PoMRI can result from Bateson–Dobzhansky–Muller (BDM) incompatibilities, in which two parental loci that have diverged cause hybrid sterility or inviability. Extending such negative epistatic interactions to the host’s microbiome adds three mechanisms that can reduce hybrid fitness: hybrid susceptibility (i.e. inferior immune systems and higher pathogen loads in hybrids), hybrid autoimmunity (i.e. upregulation of immune functions in hybrids because of negative epistasis), and cytoplasmic incompatibility (CI, i.e. hybrid embryo death caused by incompatibilities between members of the parental microbiomes)^4^. PoMRI mechanisms can operate at any developmental stage and can directly affect the fitness of both hybrid offspring and mated parental females. The best-known cases of PoMRI caused by bacteria involve *Wolbachia,* the enigmatic evolutionary omnia rotundiora comprising endosymbionts and endoparasites^5^, which infects around 40% of the world’s arthropod species^6^, with some estimates reaching 66%^7^. *Wolbachia* is famous for manipulating maternal transmission routes, and *Wolbachia* strains capable of reproductive manipulation can sweep through a population by inducing CI, or by favoring vertical maternal transmission via interference with the host’s development, resulting in parthenogenesis, feminization, or male death^8^.

Although numerous ant species are highly invasive and form allopatric populations on a global scale, nothing is known about PoMRI mechanisms involving endosymbiotic bacteria in these ecologically dominant insects. In particular, the functional role of *Wolbachia* in ant biology has thus far remained elusive, in spite of its omnipresence^9^. The globally distributed tramp ant *Cardiocondyla obscurior* has spread from its putative origin in Southeast Asia and today occurs in fruit tree plantations, city parks and other disturbed habitats in the tropics and sub tropics. In addition to *Wolbachia* populations of *C. obscurior* carry a main bacterial endosymbiont, *Cand.* Westeberhardia, which exhibits characteristics of an obligate symbiont^10^. Populations of *C obscurior*from Brazil (BR) and Japan (JP) show remarkable genomic heterogeneity and differ in a number of phenotypic traits^11^; nevertheless, mating between queens and males from the two populations readily takes place in the laboratory. However, a phenotype indicative of PoMRI was found in hybrid crosses between BR and JP populations, with BR queens mated to JP males exhibiting shorter lifespans and lower fecundity than BR queens mated to BR males^12^. At the time, this was attributed to BDM incompatibility of fast evolving sexual traits such as seminal fluid proteins, as suggested by an up-regulation of genes involved with immune response in outbred BR queens^12^.

Here, we show that the negative effect on queen fitness is in fact caused by incompatible strains of *Wolbachia* harbored by the two populations. By reducing *Wolbachia* levels in males using antibiotic treatment, we demonstrate that queen fecundity can be rescued. Genomic analyses of the *Wolbachia* strains isolated from the two populations reveal that both strains belong to *Wolbachia* supergroup A, which typically contains strains capable of reproductive manipulation. Surprisingly, CI is only induced by one of the two strains, and this may be linked to functional differences in CI-associated loci. By discovering the first case of CI between two ant populations, our study provides the first description of functional *Wolbachia* infection dynamics in ants and presents a novel perspective on the biology of globally distributed ants.

## Results

### Unidirectional hybrid incompatibility

We verified the results from^12^ by crossing queens and males from laboratory colonies of BR and JP populations collected in 2009 and 2010, respectively^11^. We conducted this experiment once in 2015 with the same outcome (Figure S1, Supplement Methods) and again in 2017/18, the results of which are presented here. As expected, compared to all other crosses the combination BR x JP (queen x male) resulted in lower fecundity as estimated by mean weekly egg number over the first six weeks after initiation of egg laying, regardless of whether the male was infected with *Cand.* Westeberhardia or not (Figure 1, Kruskal Wallis rank sum test, X^2^= 78.04, df=6, p<0.001, see Table S1 for Bonferroni-Holm corrected pairwise p-values). Similar results were obtained when maximum weekly egg numbers were compared (Figure S2, Kruskal Wallis rank sum test, X^2^=81.10, df=6, p<0.001, see Table S2 for Bonferroni-Holm corrected pairwise p-values). Strikingly, the effect was even stronger when using males naturally uninfected with *Cand.* Westeberhardia, ruling out the possibility that this endosymbiont plays a role in the induction of reproductive isolation.

**Figure 1:**
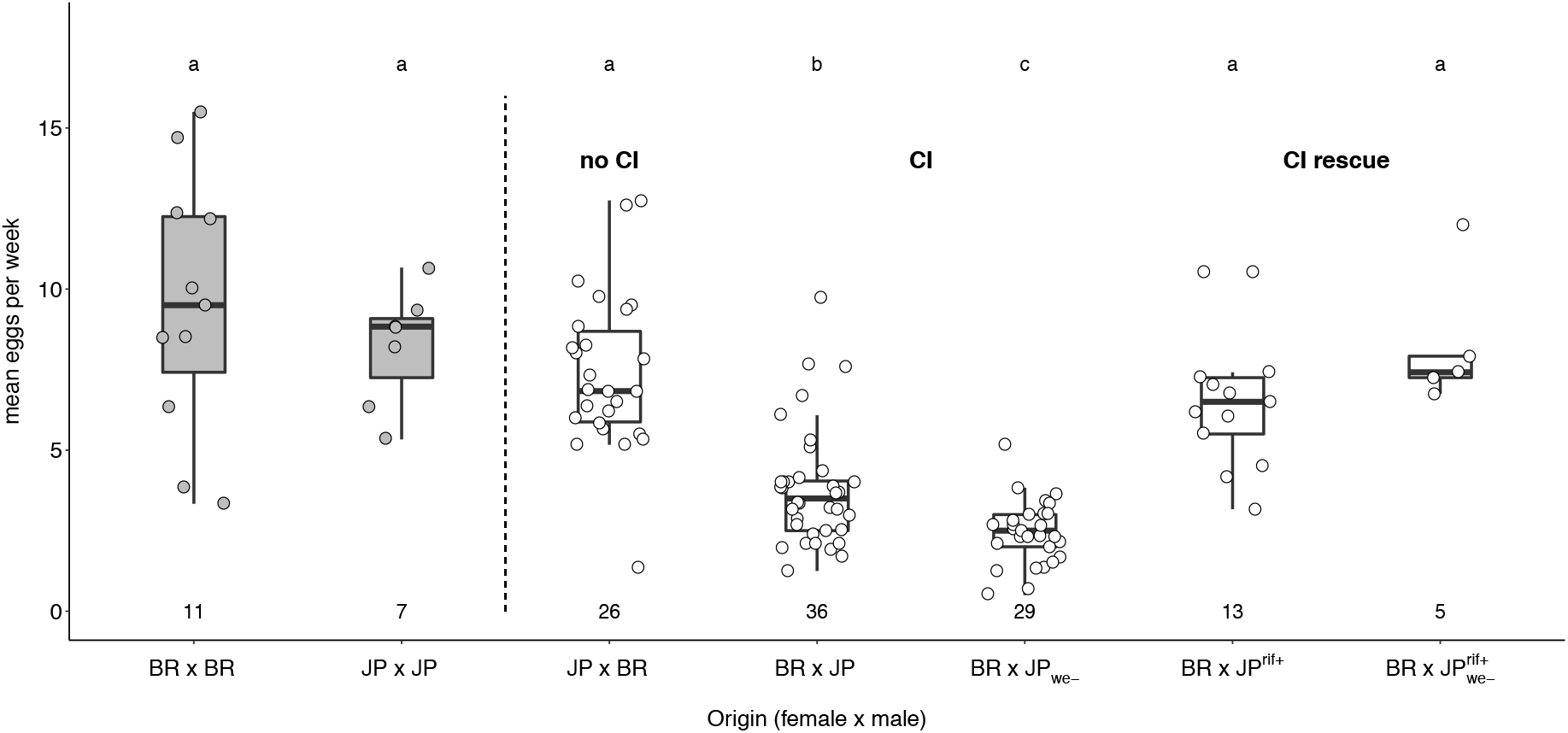
Mean weekly egg numbers produced by intra- and inter-population crosses of the ant *C. obscurior*. Mean weekly egg numbers produced by Brazilian (BR) and Japanese (JP) queens mated to males from their own or from a different population over a period of six weeks. JP ants were collected from colonies that either carried the main endosymbiont *Candidatus* Westeberhardia cardiocondylae (JP) or did not (JP_we-_). Males used in “CI rescue” crosses were collected from JP colonies with and without *Cand.* Westeberhardia cardiocondylae, which had been treated with the antibiotic rifampicin (JP^rif+^; JP_we-_^rif+^). Numbers below box plots indicate the number of replicates. Differences between groups were tested with pairwise Mann-Whitney-U-tests followed by Bonferroni-Holms correction of p-values. Letters above boxplots indicate statistically significant differences at p<0.05. For Bonferroni-Holms corrected pairwise p-values see Table S1.

We monitored a subset of inter-population crosses for an additional 6 weeks (12 weeks in total). Of the 53 BR x JP colonies that were monitored for 12 weeks, nine failed to produce diploid offspring (queen/worker pupae), and eight produced no pupae at all. We dissected the spermathecae of five of the nine male-only producing queens, and all contained sperm. All of the JP x BR colonies that were monitored for 12 weeks produced pupae, and they produced more pupae than BR x JP colonies, irrespective of whether males were infected with *Cand.* Westeberhardia (Figure 2A; Kruskal-Wallis rank sum test, X^2^=39.80, df=4, p<0.001, see Table S3 for Bonferroni-Holm corrected pairwise p-values). Furthermore, sex ratios of produced pupae were more male-biased in BR x JP colonies compared to JP x BR colonies (Figure 2B; binomial GLM: *T*_4,78_= 38.59, p<0.001, see Table S4 for Bonferroni-Holm corrected pairwise p-values), an effect caused entirely by the lack of developing females (Figure S3; Kruskal Wallis rank sum test, X^2^=5.61, df=4, p=0.23). Together, these data show consistent incompatibility between BR queens and JP males.

**Figure 2:**
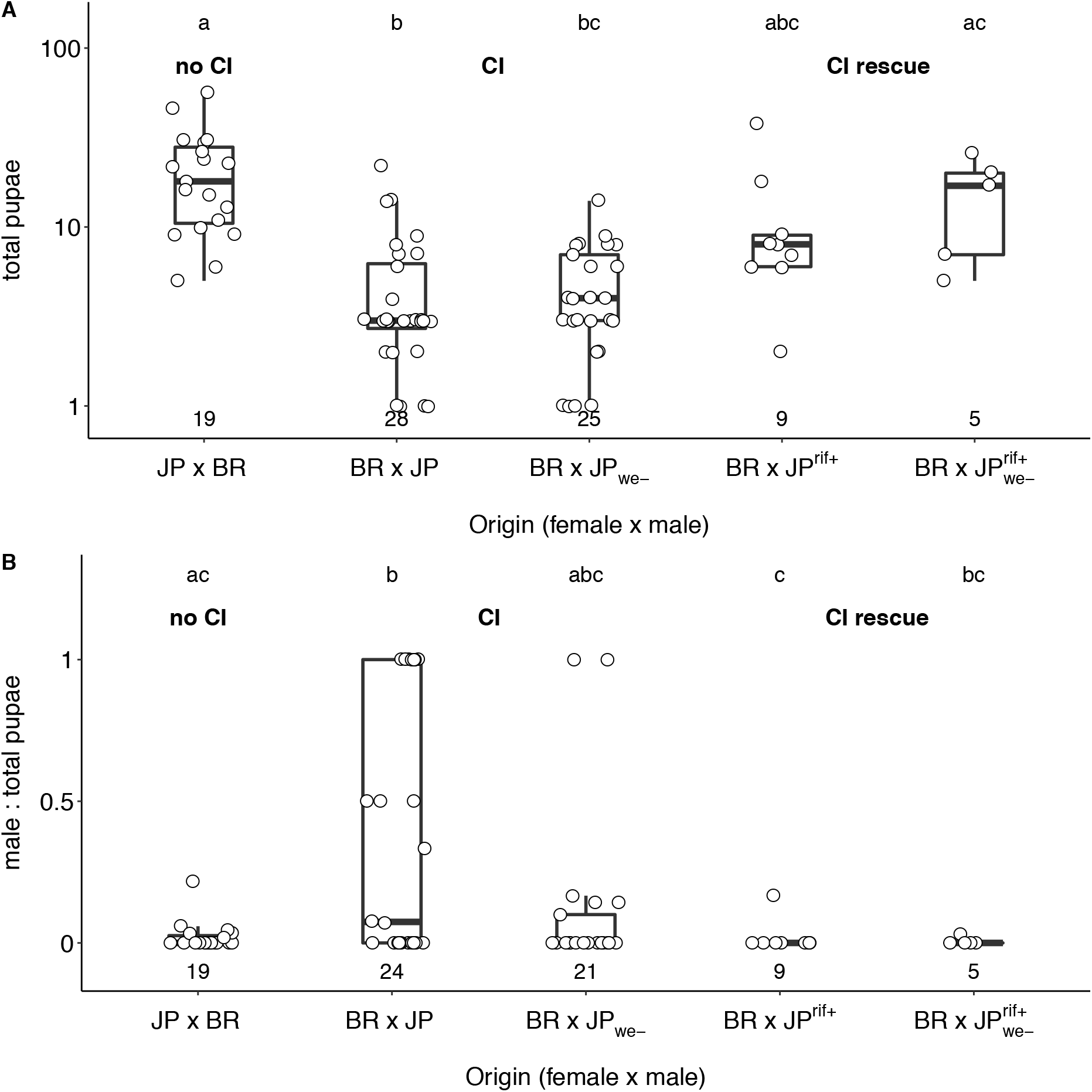
Total pupae numbers and pupae sex ratios produced by intra- and interpopulation crosses of the ant *C. obscurior*. Total number of pupae (A) and sex ratio of pupae (B) produced in crosses between Brazilian (BR) and Japanese (JP) queens and males over a period of 12 weeks. JP ants were collected from colonies that either carried the main endosymbiont *Candidatus* Westeberhardia cardiocondylae (JP) or did not (JP_we-_). Males used in “CI rescue” crosses were collected from JP colonies with and without *Cand.* Westeberhardia cardiocondylae, which had been treated with the antibiotic rifampicin (JP^rif+^; JP_we-_^rif+^). Numbers below box plots indicate the number of replicates. Total pupae numbers are plotted as log +1 for better visualization and differences between groups were tested with pairwise Mann-Whitney-U-tests followed by Bonferroni-Holms correction of p-values. For Bonferroni-Holms corrected pairwise p-values see Table S3. Sex ratios produced by different crosses were compared with a generalized linear model with logit link followed by manual Bonferroni-Holms correction of p-values. For Bonferroni-Holms corrected pairwise p-values see Table S4. Letters above boxplots indicate statistically significant differences at p<0.05.

### Cytoplasmic incompatibility

Cytoplasmic incompatibility (CI) occurs when an endosymbiont favors its transmission by making infected males incompatible with uninfected females, while infected females can mate with both infected and uninfected males. A necessary test to verify CI is ‘curing’ the host of the putative CI-causing bacteria using antibiotic treatment, which should result in the ‘rescue’ of the wild-type phenotype.

We fed JP^rif+^ colonies the antibiotic rifampicin diluted in honey every other week for 10 weeks, which resulted in the complete loss of the main endosymbiont *Cand.* Westeberhardia (Figure S4; Mann-Whitney-U-Test, W=34, df=1, p<0.001) and a significant reduction of *Wolbachia* levels in workers (Figure S4; Mann-Whitney-U-Test, W=95, df=1, p<0.01). Treating colonies with the antibiotic tetracycline did not have a negative effect on *Wolbachia* (Figure S5; Mann-Whitney-U-Test, X^2^=24, df=1, p=0.052). While *Cand.* Westeberhardia is permanently eradicated in subsequent generations the negative effect of rifampicin on *Wolbachia* is transient, with levels returning to normal 6 months after termination of treatment (Figure S6; *Cand.* Westeberhardia: Mann-Whitney-U-Test, W=64, df=1, p<0.001; *Wolbachia*: Mann-Whitney-U-Test, W=45, df=1, p=0.19). Five of 28 rifampicin-treated colonies succumbed to the treatment and died during the 10-week period, while all 28 control colonies survived.

Mating JP^rif+^ males collected shortly after ceasing antibiotic feeding with BR queens resulted in significantly rescued fecundity, although mean and maximum weekly egg numbers did not quite reach levels of BR and JP inbred queens (Figure 1, Table S1). This rescue effect was evident regardless of whether JP^rif+^ males came from *Cand.* Westeberhardia infected or uninfected colonies. Rescued BR x JP^rif+^ colonies also produced more pupae over a 12-week period than their non-rescued BR x JP counterparts (Figure 2A), and sex ratios among pupae were as female-biased as in JP x BR crosses (Figure 2B). Strongly reduced *Wolbachia* levels in JP^rif+^ males (Figure 3, Mann-Whitney-U-Test, W=163, df=1, p<0.001) support the role of *Wolbachia* in inducing hybrid incompatibility.

**Figure 3:**
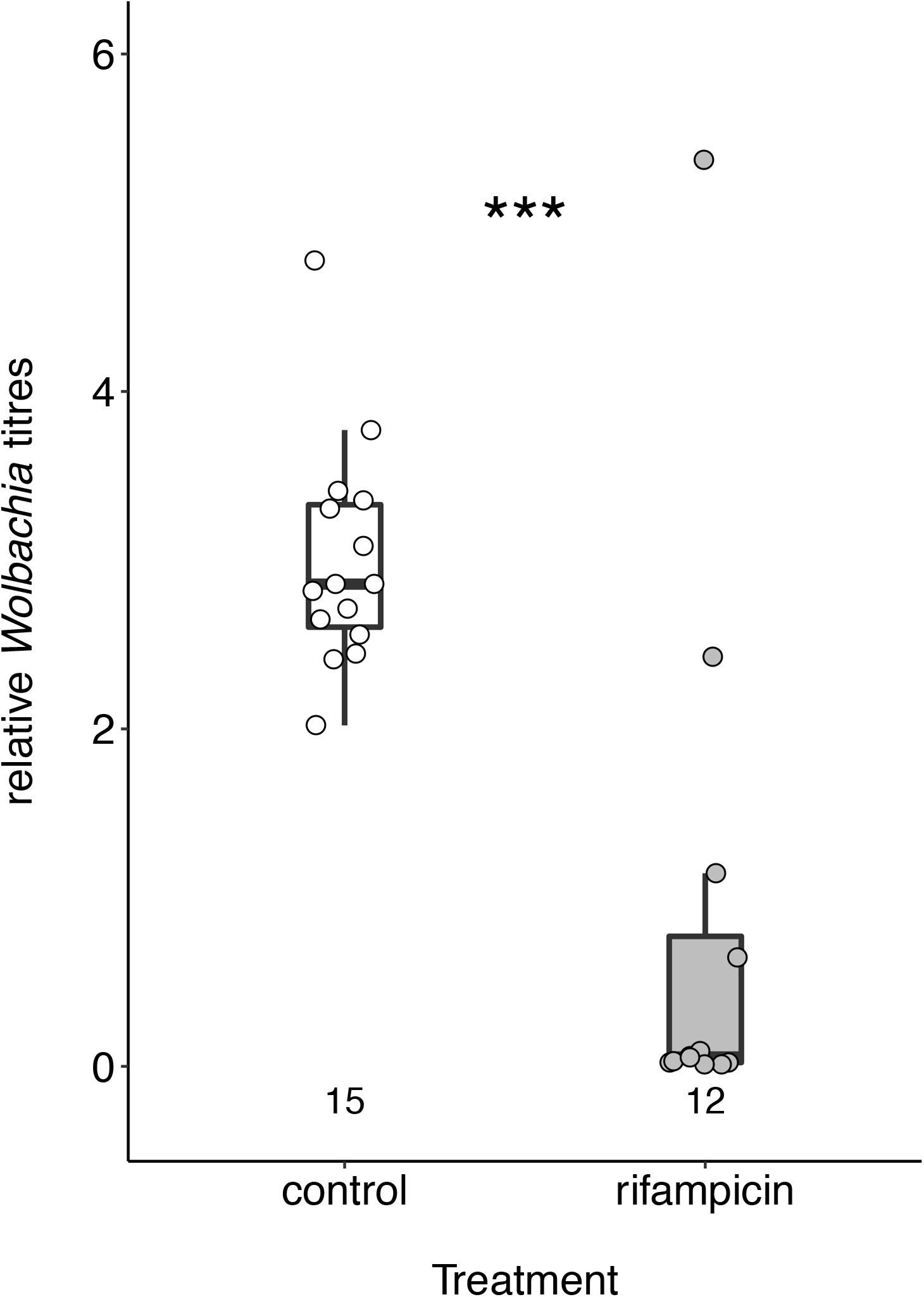
Effects of rifampicin treatment on *Wolbachia* titres in males. All males were collected from colonies that carried the main endosymbiont *Candidatus* Westeberhardia cardiocondylae. Numbers below box plots indicate the number of replicates. Differences in *Wolbachia* titres between control and rifampicin-treated males were tested with a Mann-Whitney-U-Test. Stars indicate statistically significant differences at p<0.001 (***).

We confirmed unidirectional CI with males from laboratory colonies of a second Asian population collected in Taiwan (TW), which is infected with a *Wolbachia* strain that amplifies a *wsp* sequence (coding for the hypervariable *Wolbachia surface protein)* identical to those from JP samples (Figure S7). We set up crosses with queens from BR (BR x TW) and JP (JP x TW), and the reciprocal combination (TW x JP). Again, only the combination BR x TW resulted in reduced fecundity, both when mean weekly egg numbers (Figure S8; Kruskal Wallis rank sum test, X^2^=9.96, df=2, p<0.01, see Table S5 for Bonferroni-Holm corrected pairwise p-values) and maximum weekly egg numbers over six weeks (Figure S8; Kruskal Wallis rank sum test, X^2^=10.19, df=2, p<0.01, see Table S6 for Bonferroni-Holm corrected pairwise p-values) were compared.

### *Wolbachia* genomics and phylogeny

We assembled the *Wolbachia* genomes from BR (*w*Cobs-BR) and JP (*w*Cobs-JP) populations of *C. obscurior* using Illumina HiSeq2000 100-bp reads of 200 and 500-bp insert paired-end libraries^11^. The genomes of both *Wolbachia* were assembled to 148 (BR) and 182 (JP) scaffolds (Table 1). Both strains showed similar genome sizes and G + C contents, with an estimated 1,013 (BR) and 1,058 (JP) predicted protein-coding genes. An analysis on shared genes revealed that both strains share a core of 907 proteins, with 85 unique to *w*Cobs-BR and 109 unique to *w*Cobs-JP. According to Bayesian phylogenetic analyses of ribosomal proteins from 23 *Wolbachia* strains, both *Wolbachia* strains belong to supergroup A, which together with supergroup B contains strains capable of reproductive manipulation. *w*Cobs-JP was found to be more closely related to *w*Ha and *w*CauA strains, while *w*Cobs-BR was phylogenetically closer to *w*Au and *w*Mel (Figure 4A). A synteny analysis revealed that both *w*Cobs-JP and *w*Cobs-BR strains keep general synteny, but with several rearrangements (Figure 4B). This is consistent with previous observations of group A *Wolbachia* strains^13^.

**Figure 4:**
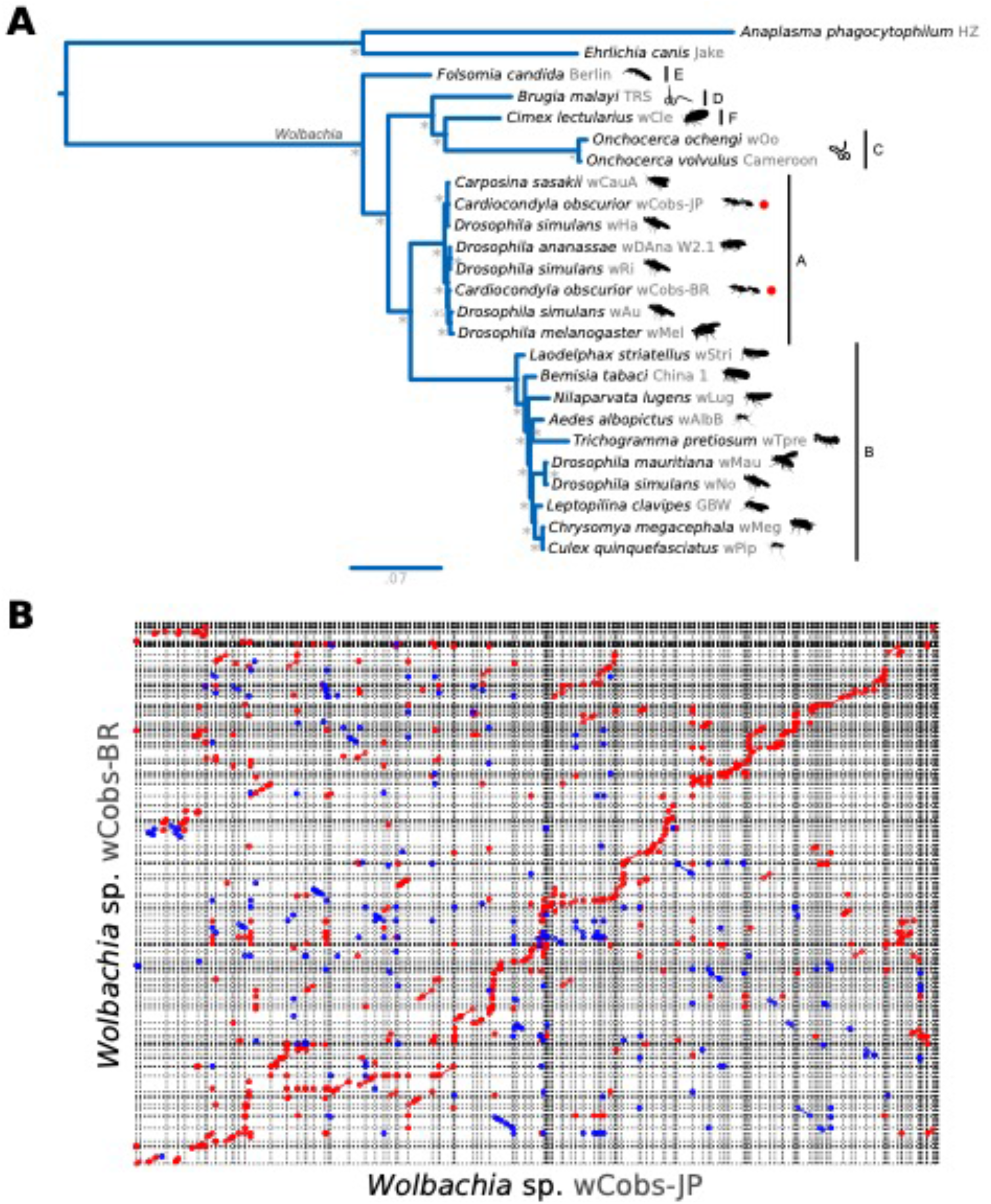
*Wolbachia* spp. phylogeny and genome synteny. (A) Bayesian phylogenetic placement of *Wolbachia* strains sequenced from *C. obscurior* (*w*Cobs-BR and *w*Cobs-JP). *Anaplasma phagocytophilum* and *Ehrlichia canis* were used as outgroups. For *Wolbachia,* names at tips correspond to the host from which they were sequenced/isolated. Red dots are used to highlight the newly sequenced strains. Next to the binomial names, strain names are shown in grey and silhouettes for the host are sketched in black. Horizontal bars delimit *Wolbachia* supergroups (A-F). *=1.0 posterior probability. (B) Genome-wise nucleotide-based synteny plot between *w*Cobs-BR and *w*Cobs-JP. Red is used to indicate a direct match and blue a reverse complement match.

**Table 1:** Wolbachia genome assembly statistics. Genome assembly and draft annotation statistics for the newly sequenced *Wolbachia* strains from *C. obscurior* (*w*Cobs-BR and *w*Cobs-JP).

|  | Wolbachia |  |
| --- | --- | --- |
|  | wCobs-BR | wCobs-JP |
| **Assembled genome size (Mbp)** | 1.19 | 1.30 |
| **Number of scaffolds** | 148 | 182 |
| **N50** | 14,698 | 16,173 |
| **G+C content (%)** | 35.3 | 35.1 |
| **Predicted CDSs\*** | 1,013 | 1,058 |
| **tRNAs** | 34 | 34 |
| **rRNAs** | 3 | 3 |
| **Other ncRNAs genes/motifs** | 71 | 54 |

### *Wolbachia* phenology

To compare infection titers between populations we performed qPCR of the *Wolbachia* gene *Cytochrome c oxidase subunit 1 (coxA)* against the ant housekeeping gene *elongation factor 1-alpha 1 (EF1).* JP queens had higher relative *w*Cobs-JP titers than BR queens infected with *w*Cobs-BR (Figure S9, Mann-Whitney-U-Test, W=23, df=1, p=0.043). Likewise, JP workers showed higher infection titers with *w*Cobs-JP compared to BR workers infected with *w*Cobs-BR (Figure S9, Mann-Whitney-U-Test, W=16, df=1, p=0.017).

To test for morph-bias in infection we compared *coxA* expression in adult queens, workers, winged males and wingless males using qPCR. *coxA* levels did not differ between sexes and were consistently higher in winged morphs (queens, winged males) compared to wingless morphs (workers, wingless males) (Figure S10, Kruskal Wallis rank sum test, X^2^=19.53, df=3, p<0.001).

To measure the potential cost of *Wolbachia* infection and to test for a selective loss in workers, we compared infection levels of workers and queens over time. In contrast to *Cand.* Westeberhardia^10^, *Wolbachia* levels quickly reached a plateau in both queens and workers. There was no indication that individuals lose *Wolbachia* over the first few weeks of their lives, which typically last for up to three (workers) or six months (queens) (Figure S11, workers: ANOVA, F_4,26_= 2.69, p=0.059; queens: ANOVA, F_5,50_= 15.95, p<0.001).

### CI mechanism

CI usually results in the failure to produce viable zygotes but how CI operates functionally is still not clear. The two championed models are based on 1) a sperm modification step during spermatogenesis which is reversed by a rescue factor or 2) the transfer of a toxic product via the sperm and subsequent binding to a female-derived antidote^14–17^. The current genetic view for CI in *Drosophila* favors a two-by-one model^17^, involving two adjacent genes (coined *cifA* and *cifB)* located in a region of prophage origin in i/vMel^16–21^, which contains several genes with eukaryote function^22^. Expression of both *cifA* and *cifB* in males is required to cause CI, while expression of *cifA* rescues embryo mortality in females^21^. Phylogenetic analyses of the Cif proteins have revealed at least 5 distinct phylogenetic clades referred to as Types 1 −5, respectively^18,23,24^.

We found homologues of *cifA* and *cifB* in *Wolbachia* strains from BR and JP. In the latter, we found two additional homologues. Through Bayesian phylogenetic inference, and following the classification scheme of^23^, we found that all copies of *cifA* and *cifB* belong to Type I (Figure 5A, B). The *cifA* and *cifB* genes found in NODE_002 of *w*Cobs-JP are closely related to those found in *w*Cobs-BR and in the cryptic phage of the strain uvSol isolated from the fig wasp *Ceratosolen solmsi* (Figure 5A, B). The *cifA* and *cifB* genes found in NODE_063 of *w*Cobs-JP show a more distant relationship to their homologues in *w*Cobs-BR. Following^23^, we performed a protein remote homology search with the webserver of HHpred’s v3.2.0 webserver^25^. While the *cifA* gene of *w*Cobs-BR contained a DUF249 domain, neither *cifA* homologues in *w*Cobs-JP showed homology to known domains (Figure 5C). In contrast, putative domains were found in all *cifB* genes. These shared a putative nuclease domain (PD-(D/E)XK nuclease/DpnII-MboI) found in other type I *cifB* genes. The *cifB* homologue found in NODE_063 in *w*Cobs-JP shared most domains with the *cifB* homologue in *w*Cobs-BR, with an additional peptidase domain (Peptidase_C58; PF03543).

**Figure 5.**
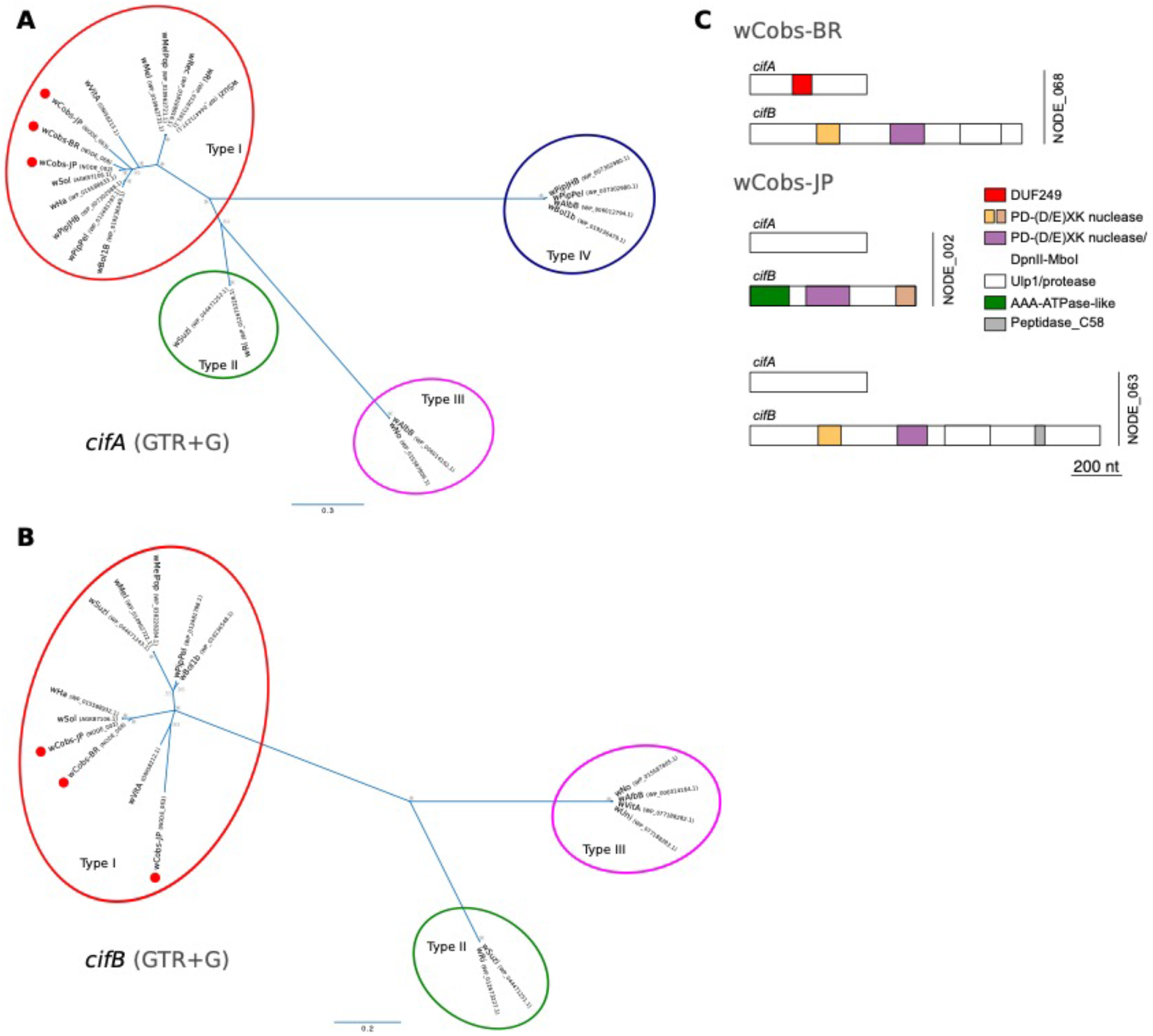
Phylogeny and domain structure of *cif* genes. Bayesian phylogenetic trees for (A) *cifA* and (B) *cifB* genes. Colored ovals are used to highlight the different gene “types”. *= 1.0 posterior probability. (C) Domain structure for the *cif*genes of *w*Cobs-BR (top) and *w*Cobs-JP (bottom) strains. For the latter, the two loci are shown and indicated with a vertical bar to the right of the sketches. Color codes and designation of the domains are shown in the legend.

### *Wolbachia* in males

As mentioned above, *Wolbachia*-induced CI has often been linked to sperm modification during spermatogenesis^26^ or the transfer of a CI toxin along with or in the sperm. The presence of *Wolbachia* in the testes is thus expected, although their specific localization in spermatocytes or spermatids might not be a prerequisite for inducing CI, as has been observed in the wasp *Nasonia vitripennis*^27^. We used fluorescence in situ hybridization to localize *Wolbachia* in the abdomens of male *C. obscurior,* confirming the presence of the bacteria throughout different tissues including the testes in both JP (Fig. 6A-C) and BR (Figure 6D) individuals. *Wolbachia* was also observed co-infecting bacteriocytes dominated by *Cand.* Westeberhardia, yet in significantly lower densities (Figure 6C).

**Figure 6:**
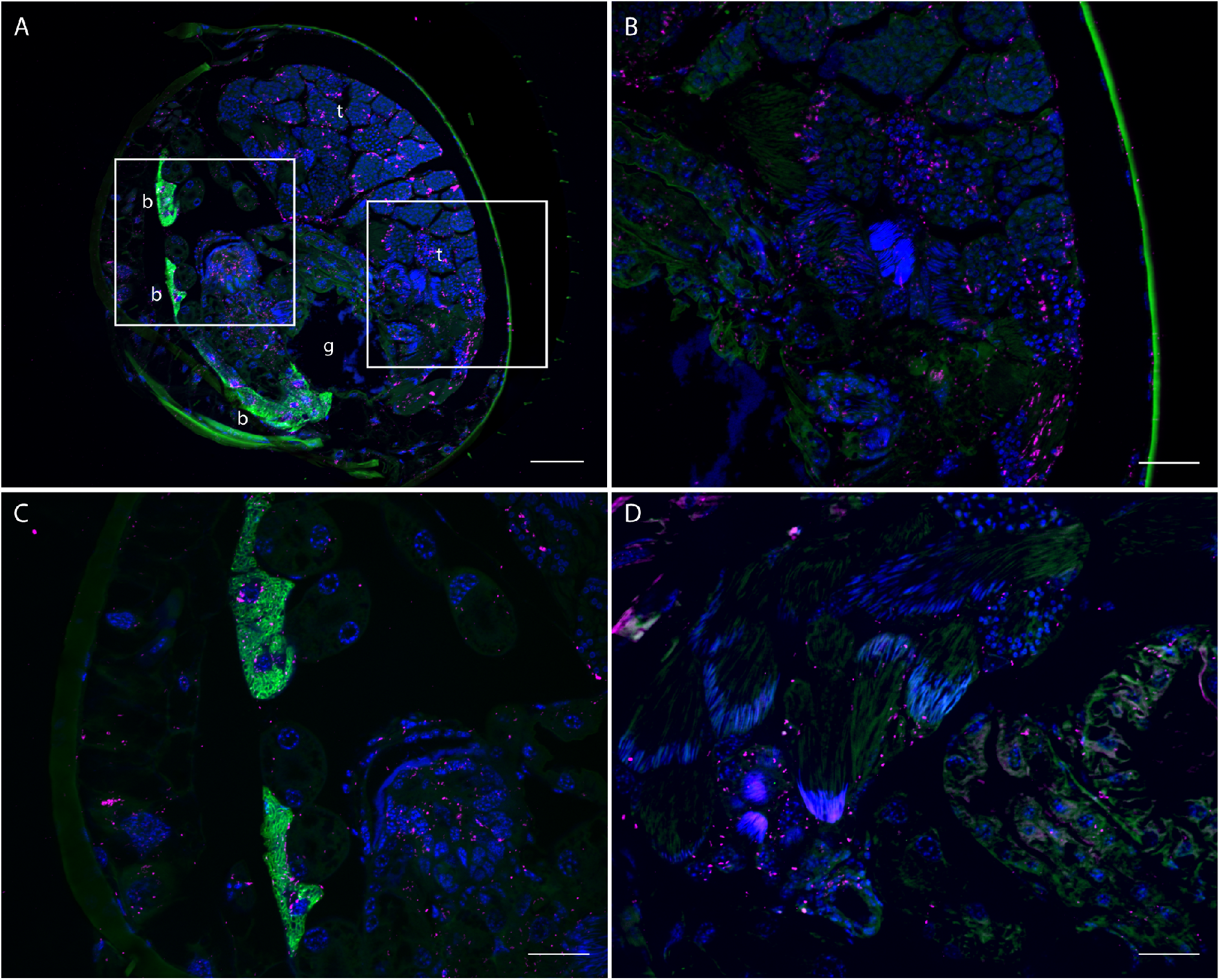
Localization of endosymbiotic bacteria in male adult abdomens of *C. obscurior* via fluorescence in situ hybridization. *Wolbachia* were specifically stained with two Cy5-labeled probes (magenta), and *Cand.* Westeberhardia with a Cy3-labeled probe (green). Host cell nuclei are counterstained with DAPI (blue). (A) Transversal section of *C. obscurior* JP male abdomen showing the testes (t), the *Cand.* Westeberhardia-containing bacteriomes (B) and the gut (g). The areas indicated by rectangles are displayed in higher magnification showing *Wolbachia*-infected seminiferous tubules (B) and bacteriomes co-infected with *Cand.* Westeberhardia and *Wolbachia* (C). (D) *C. obscurior* BR male testes and gut epithelium infected with *Wolbachia.* Scale bars: 50 μm (A) and 20 μm (B-D).

## Discussion

Insect-*Wolbachia* interactions can range from beneficial to detrimental and from facultative associations to rare cases of ultimate mutualisms where hosts have higher fitness with *Wolbachia* than without^5^. This prevalence and diversity of *Wolbachia* associations predictably also occurs in ants; however, our knowledge of ant-*Wolbachia* interactions is limited^9^. In the few ant-*Wolbachia* associations studied so far, infection is facultative (e.g.^28–30^), indicating ongoing arms races between hosts and symbionts^31^. One particularly obvious question to ask is whether *Wolbachia* has the power to control sex allocation, a trait shaped by kin selection in social Hymenoptera^32^. In this context, facultative *Wolbachia* infections have been reported to cause transient changes in sex allocation^29^, or to have no effect^33^. Similarly contrasting results have been obtained regarding effects on colony productivity^29,34^. It has furthermore been speculated that *Wolbachia* can bias caste ratios towards the production of queens^28,35^, but current data are scarce and provide little support for this hypothesis^9^. By providing the first functional description of an ant-*Wolbachia* association involving two strains of *Wolbachia*, our study lays the groundwork for in-depth investigations into how *Wolbachia* affect ant reproductive biology and fitness.

*Wolbachia* clearly play a significant role in the biology of *C. obscurior*. First, all screened individuals were infected regardless of population, caste or sex. *Wolbachia* titers were higher in queens and winged males compared to workers and wingless males, a difference that is likely associated with the relative amounts of reproductive tissue as *C. obscurior* workers lack ovaries and winged males have larger testes than wingless males^36^. Second, rifampicin treatment merely reduced *Wolbachia* titers, and this effect was reversible. This was not the case for *Cand.* Westeberhardia, which was permanently cleared by rifampicin treatment. Third, *Wolbachia* titers did not decline with age in workers. *Wolbachia* infection can be costly and selection may thus favor reduced infection loads in sterile workers, as these do not contribute to vertical transmission^28,35^. Such a mechanism appears to be acting on *Cand.* Westeberhardia, where titers in workers (but not queens) decreased significantly with age, indicating that while the host may benefit from infection during development or in the adult egg-laying queen stage, the symbiont is not required^10^. Consistently high titers in workers indicate that *Wolbachia* do not have deleterious effects on worker performance in *C. obscurior*, unlike *Wolbachia* in *Acromyrmex* ants^37^. To what degree the population-specific strains differ in their effect on host phenotype remains unclear. Higher *Wolbachia* titers in JP individuals suggest that the *w*Cobs-JP strain may be better integrated into the host’s biology. Alternatively, these differences may be driven by population-specific competition with *Cand.* Westeberhardia over limited resources.

Sex allocation in *C. obscurior* is female-biased^38^ and queens presumably have full control over male production, as investment into males increases in the presence of competing queens^39–41^, Accordingly, we found no indication that *Wolbachia* influences sex allocation. Queens from non-CI (JP x BR) crosses produced few males in the first 12 weeks of their lives. This may be a consequence of the short observation period, equaling ~50% of the average queen lifespan^38^. The sex ratios produced by CI (BR x JP) crosses showed a bimodal distribution, with several queens matching sex ratios produced by non-CI queens (albeit with lower overall productivity), while others produced only males. On average, however, CI queens did not produce more males than non-CI queens. There was also no indication that *Wolbachia* altered caste ratios towards production of queens (data not shown). Concordantly, a previous study comparing investment into sexuals and workers did not find differences between the BR and JP populations^38^. Testing directly for a caste-biasing effect of *Wolbachia* requires complete elimination of the bacterium. While this cannot be done without also killing the ant host, preliminary trials indicate that cross-infection of the two *Wolbachia* strains is possible. In the future, such manipulations may help clarify whether *Wolbachia* have caste-manipulating power in *C. obscurior*.

Global exchange of stowaway ants leading to secondary contact between allopatric populations is common (e.g. in the tropical fire ant *Solenopsis geminata*^42^, and the little fire ant *Wasmannia auropunctata*^43^) and thus the presented scenario, albeit conceived in the laboratory, is not an unrealistic one. A previous study emphasized the role of BDM incompatibility between allopatric *C. obscurior* populations in causing a CI-like phenotype^12^. *Wolbachia*-induced incompatibilities were ruled out because of identity of a partial sequence of *wsp* extracted from seven individuals (three from BR colonies collected in 2004, four from JP colonies collected in 2005) using the wsp81F/wsp691R primer combination^44^. Gene expression comparisons of allopatrically and sympatrically mated BR queens pointed towards elevated immune responses and other changes that mimicked a CI-like phenotype. In *Drosophila, Wolbachia* can manipulate expression of key regulatory genes during spermatogenesis, which leads to embryonic lethality and a CI-like phenotype^45^. Thus, what appeared to be BDM incompatibility may have been caused by *Wolbachia* inducing a CI-like phenotype via interference with queen gene expression. The alternative and more parsimonious explanation for the CI-like phenotype based on the results from the present study is a classic unidirectional *Wolbachia*-driven CI effect involving incompatible *Wolbachia* strains. This also explains why JP x BR crosses did not elicit a CI phenotype. The overexpression of immunity-related genes in allopatrically mated BR queens detected by Schrempf et al. 2015 may simply reflect physiological responses to aborted oogenesis, as opposed to responses to ejaculate quality, or to injuries caused by non-matching male genitalia. Furthermore, we recently found that some BR colonies were infected with an additional *Wolbachia* strain displaying partial *wsp* sequences identical to those of *w*Cobs-JP (data not shown), indicating horizontal transfection in our rearing facilities, especially in older laboratory colonies.

The mechanism of CI in *C. obscurior* appears to align along the *Wolbachia cif* gene axis described in *Drosophila* fruit flies and *Culex* mosquitoes^15,17,19,23,46^. This genetic region, which is of prophage origin and capable of manipulating the eukaryote host^18^, showcases the importance of using a hologenomic approach to study evolution^47^. Similar prophage-derived regions containing pairs of homolog *cif/cif*-like genes have since been described in a number of *Wolbachia* strains^23,24^. Surprisingly, the homologs of *cifB* in *w*Pip and *w*Mel have different biochemical functions: both putatively contain ankyrin-like repeats for protein-protein interaction with eukaryotic DNA^46^ but *cifB*_*w*Mel_ is not a putative deubiquitinase^16^, as postulated for *cifB*_*w*Pip_^19^. Both *Wolbachia* strains isolated from *C. obscurior* have homologs of *cifA* and *cifB* but only the *w*Cobs-JP strain has an additional *cifA*/*B* pair, pointing towards a role for the latter pair in inducing reproductive isolation. Whether the second *cifA*/*B* containing region was lost in *w*Cobs-BR, or whether it stems from a duplication event and subsequent divergence in *w*Cobs-JP as has been described for the *w*AlbB strain of *Aedes albopictus*^48^, remains to be shown. Alternatively, functional differences in *cifA/B* homologs between *w*Cobs-JP and *w*Cobs-BR, or an interaction of the two *cif* gene products in JP males could be causing CI. Irrespective of the exact CI mechanism, we do not know why a mere reduction of *Wolbachia* titers in males was sufficient to rescue queen fecundity. This requires studying the effects of *Wolbachia* and antibiotic treatment on sperm production and development. One possibility is that rifampicin treatment causes ribosomal damage that prevents *Wolbachia* from modifying sperm. At the same time, antibiotic-induced damage to sperm mitochondria^49^ may explain the slightly lower productivity of “CI rescue” crosses compared to non-CI crosses. It is also unclear why the CI effect was stronger on egg numbers but not pupae numbers when using males from colonies naturally uninfected with *Cand.* Westeberhardia, but this could point toward interaction effects between the two symbionts.

Our study raises many questions about the evolutionary history of the *C. obscurior-Wolbachia* association. The putative origin of *C. obscurior* in Southeast Asia, together with sequence identity between *Wolbachia* strains isolated from populations found in Japan and Taiwan, suggest that these strains represent the ancestral association. Geographic patterning separating New from Old World *Wolbachia* strains has been observed previously in ants^50^, and frequent horizontal transmissions^51,52^ can lead to a surprising diversity within species^53^. *C. obscurior* and its sister species *C. wroughtonii* are the only known representatives of the 100+ described species in the genus *Cardiocondyla* (the so-called heart-node ants) that have adopted an arboreal lifestyle, as opposed to nesting in soil and rock crevices. This makes them highly susceptible to horizontal transmission of *Wolbachia* from other plant-associated insects. Although *Wolbachia* from different supergroups may co-exist in a host^13^, it is unlikely that two strains from the same supergroup co-exist without eventually recombining and merging. Generally, unidirectional CI should facilitate introgression between populations, while bidirectional CI should promote divergence. This predicts mixed *C. obscurior* genotypes but only the presence of the *w*Cobs-JP strain in secondary contact zones, assuming that hybrids perform as well as parental lineages. How then has the ancestral Asian strain been replaced? *Wolbachia* acquisition and persistence in *C. obscurior* may be driven by selection on the strain that is better adapted to the local environment, perhaps analogous to *Wolbachia*-driven thermal preferences in *Drosophila* flies^54,55^. Resolving how *w*Cobs-BR replaced the putatively ancestral *w*Cobs-JP strain, which holds the power of CI, requires testing how mixed combinations of host and *Wolbachia* perform under varying ecological conditions. Coupled with a global survey of *Wolbachia-C. obscurior* associations, this will help unravel how the symbiosis shapes the worldwide distribution of this tramp ant.

## Conclusion

One important consequence of unidirectional reproductive isolation is unequal inheritance, which allows a BDM-causing locus or a reproductive manipulator to quickly spread in secondary contact zones. Such disruptive factors can be more powerful than natural selection on other fitness-relevant traits, and lead to the evolution of PoMRI and hybrid inviability, processes estimated to require several million years in *Drosophila* flies and *Pogonomyrmex* ants^56,57^. Our discovery of reproductive isolation between allopatric ant populations, driven by two naturally occurring, closely related *Wolbachia* strains that differ in their genomic makeup, demonstrates that these evolutionary processes can be observed in real time and lays the groundwork for detailed studies of *Wolbachia*-ant biology. In particular, a system in which CI is accompanied by a CI-like phenotype in host gene expression may prove useful for understanding the function of the *cif* gene family. More generally, hologenomic approaches such as the one presented here will help tackle the challenges associated with human-induced changes to ecological and evolutionary dynamics.

## Methods

*Cardiocondyla obscurior* presumably originates from Southeast Asia and is now distributed around the tropics and subtropics. Queens in *C. obscurior* are produced year-round and have a short generation time of ~120 days (from egg to queen to egg). Colonies usually contain 2-5 queens^11^ and a single wingless male that can live up to a few months and continually produces sperm^58^. Such males transfer on average 1000 sperm cells per copulation^59^, which is sufficient for a lifelong production of diploid offspring by an inseminated queen. Newly hatched queens can decide to stay in the maternal nest or to leave - alone or with a small group of workers - irrespective of mating status. Like all Hymenoptera, ants exhibit haplodiploid sex determination and virgin females can produce males from haploid, unfertilized eggs. Virgin queens in *C. obscurior* can copulate with their own sons (pers. obs.), enabling the production of workers and queens, similar to a related *Cardiocondyla* species^60^. This biology has facilitated the worldwide spread of several species of the genus^61^.

In this study, we used inbred laboratory ‘populations’ derived from colonies collected in Bahia, Brazil (BR) in 2009 (Brazilian Ministry of Science and Technology, permits 20324-1/40101-1) and Okinawa, Japan (JP) in 2010. The JP population exhibits a natural polymorphism regarding the presence of the endosymbiont *Candidatus* Westeberhardia cardiocondylae, with some colonies being naturally uninfected (JP_we-_)^10^. Although the BR and JP populations differ in cuticular hydrocarbon composition, queen size and behavior^11^, workers are morphologically indiscernible according to exploratory data analyses by hierarchical and nonhierarcical, vector-based forms of Nest Centroid clustering: NC-Ward, NC-part.hclust, NC-part.kmeans, NC-NMDS-kmeans^62,63^. These analyses considered 15 continuous morphometric characters. Specifically, when considering 82 cosmopolitan nest samples of the sister taxa *C. wroughtonii* and *C. obscurior*, all JP and BR samples were allocated to the latter species by all four types of NC-clustering. This result was confirmed with posterior probabilities of p > 0.98 if these samples were run as wild-cards in a linear discriminant analysis. Analyses of 32 nest samples representing only the *C. obscurior* cluster could not allocate the JP and BR populations to different intraspecific clusters nor could these analyses demonstrate any intraspecific structuring at all. This suggests that *Wolbachia*-induced reproductive isolation has not yet led to a divergence into morphospecies. In addition to JP and BR colonies, we used colonies collected from a second Asian population in Taipei, Taiwan in 2013 (TW) to confirm the presence of unidirectional CI. For all experiments, colonies were kept in square plaster-bottom nests (100 x 100 x 20 mm, Sarstedt, Germany) and held in climate chambers under a 12h/12h light/dark cycle and a 28°C/22°C temperature cycle. Food (honey and *Drosophila* or pieces of *Periplaneta americana*) and water were provided to stock colonies every 3 days. All animal treatment guidelines applicable to ants under international and German law were followed. Collection of colonies that form the basis of the laboratory population used in this study was permitted by the Brazilian Ministry of Science and Technology (RMX 004/02). No other permits were required for this study.

## Antibiotic treatment

Rifampicin is a broad-spectrum antibiotic that acts via inhibition of bacterial RNA polymerase^64^. To produce *Wolbachia*-cured males, we split large *Cand*. Westeberhardia-infected (JP) and uninfected (JP_we-_) stock colonies into two equal halves. Half of these splits were designated as controls (JP: n = 13, JP_we-_: n=5), while the other half was treated with antibiotics (JP: n=13, JP_we-_: n=5). We diluted 0.0025 g of solid rifampicin (Sigma-Aldrich, USA) in 500 μl of a 1:1 honey-water solution for a final antibiotic concentration of 0.5%. The antibiotic solution was placed on a shaker for 4 hours to ensure complete mixing, and subsequently covered with aluminum foil and stored at −20°C. Treated colonies were fed with the antibiotic solution twice per week every other week, for a total duration of ten weeks. On the days following treatment, the antibiotic solution was removed from the nest. Between antibiotic feedings (i.e. every other week), colonies were fed with honey and pieces of autoclaved cockroach (to prevent bacterial re-infections) and water ad libitum. Control colonies were fed twice a week with honey and cockroaches and received water ad libitum.

### *Wolbachia* and *Cand*. Westeberhardia infection

We assessed the presence and titers of bacterial infections using PCR and qPCR, respectively, on DNA extracted from whole bodies (worker pupae & adults, queen pupae & adults, adult males). DNA was extracted from samples using a standard CTAB protocol.

### Presence of infection

We confirmed infection in a subset of workers from BR and JP colonies. Infection with *Wolbachia* was tested with a PCR assay using primers specific for a fragment of the *Wolbachia surface protein* gene of the BR (105-bp, *wsp_wCobs_BR-for: 5’- ‘TAAATCTTGCATCTGTAACATT-3’, wsp_wCobs_BR-rev: 5’- CTGCGGATACTGATACAACTACTG-3’)* and JP (254-bp, *wsp_wCobs_JP-for: 5-CATTTTGACTACTCACAGCGGTTG-3’, wsp_wCobs_JP-rev: 5’CTGCGGATACTGATACAACTACTG-3’)* strains. *Cand.* Westeberhardia infection was tested by amplifying a 204-bp fragment of the *ribonucleoside-diphosphate reductase 1 subunit beta* gene of *Westeberhardia (WEOB_403) (nrdB_weCobs-for:* 5’-GGAAGGAGTCCTAATGTTGCG-3’; nrdB_*weC*o*bs*-rev: 5’-ACC AGAAATATCTTTTGCACGTT-3’). A 104-bp fragment of the *C. obscurior* gene *elongation factor 1-alpha 1 (Cobs_01649) (EF1-for:* 5’-TCACTGGTACCTCGCAAGCCGA-3’; *EF1-rev:* 5’-AGCGTGCTCACGAGTTTGTCCG-3’) was used as a positive control. In addition, infection was assessed in two adult queens from each of five TW colonies to confirm that this population carries the *w*Cobs-JP strain. Each PCR reaction contained 5 μL Taq polymerase (GoTaq, Promega), 3 μL H_2_O, 0.5 μL forward primer (10 μM), 0.5 μL reverse primer (10 μM) and 1 μL DNA. PCRs were run at 94°C for 4 minutes followed by 36 cycles at 94°C for 30s, 57°C for 30s and 72° for 30s, with a final step of 72°C for 10 minutes. Electrophoresis separation of PCR products was performed on a 1.5% TAE-Gel with 0.5 μl of 0.001 % Gel-Red for 45 minutes at 60 V and 65 mA.

### Population-specific Wolbachia titers

We compared *Wolbachia* infection titers of adult workers and queens from BR (n=9 workers, n = 10 queens) and JP (n = 10 workers, n=9 queens) colonies by amplifying a portion of the *Cytochrome c oxidase subunit 1* gene *(coxA_wCobs-for.* 5’-TTGGTCATCCAGAAGTTTACGT-3’, *coxA_wCobs-rev.* 5’-TGAGCCCAAACCATAAAGCC-3’) in qPCR. As a relative standard *elongation factor 1-alpha 1 (Cobs_01649)* was used. Each qPCR reaction contained 5 μL SYBR (Peqlab), 2 μL H_2_O, 1 μL forward primer (2 μM), 1 μL reverse primer (2 μM) and 1 μL DNA. Reactions were performed in duplicates and run at 95°C for 3 minutes followed by 40 cycles at 95°C for 5s, 60°C for 20s, and 95° for 1 0s, and a final melt curve analysis step in which reactions were heated from 65°C to 95°C in 5°C steps. Single-amplicon production was confirmed with melt curve analyses and relative *Wolbachia* titers calculated with the 2^-ΔCT^ method^65^.

### Age and morph-specific Wolbachia titers

We quantified *Wolbachia* titers in individual queens and workers from BR colonies by determining relative *coxA*_*wCobs* copy numbers with qPCR across developmental stages and adult age classes. In workers, titers were measured in pupae (n=4 white pupae, n=5 dark pupae) and in adults two days (n=5), 14 days (n=7), and 28 days (n=6) post-hatching. In queens, titers were measured in pupae (n=5 white pupae, n=4 dark pupae), as well as in adults two days (n=7), 14 days (n=7), 28 days (n=10 virgin queens, n=7 mated queens) and 48 days (n=9) post-hatching. In addition, we quantified *coxA_wCobs* copy numbers in adult morphs (n = 15 queens, n = 12 workers, n=6 winged males, n=7 wingless males). For all reactions, *elongation factor 1-alpha 1 (Cobs_01649)* was used as a housekeeper. Each qPCR reaction contained 5 μL SYBR (Peqlab), 2 μL H_2_O, 1 μL forward primer (2 μM), 1 μL reverse primer (2 μM) and 1 μL DNA. Reactions were performed in triplicates and run as described above. Single-amplicon production was confirmed with melt curve analyses and relative quantities of *Wolbachia* calculated with the 2^-ΔCT^ method.

### Rifampicin effects on infection titers

We verified the efficacy of rifampicin treatment by measuring *Wolbachia* and *Westeberhardia* titers in dark worker pupae produced by antibiotic-treated (n=21 from 9 colonies) and control colonies (n=21 from 8 colonies) using qPCR of *coxA*_*wCobs and nrdB_weCobs*. In addition, we measured the *Wolbachia* titers of JP males used in mating combinations BR x JP and BR x JP^rif+^ (JP: n=15, JP^rif+^: n=12) with qPCR of a portion of the JP-specific *wsp* gene *(wsp_wCobs_JP).* For all samples, *elongation factor 1-alpha 1* (*Cobs_01649*) was used as a housekeeper. Each qPCR reaction contained 5 μL SYBR (Peqlab), 2 μL H2O, 1 μL forward primer (2 μM), 1 μL reverse primer (2 μM) and 1 μL DNA. Reactions were performed in duplicates (males) or triplicates (worker pupae) and run as described above (except with an annealing temperature of 57°C for male pupae). Single-amplicon production was confirmed with melt curve analyses and relative quantities of *Wolbachia* and *Westeberhardia* calculated with the 2^-ΔCT^ method.

### Experimental crosses

We designed seven mating combinations between queens and males from the two populations: 1) JP x JP (n = 7), 2) JP x BR (n=26), 3) BR x BR (n=11), 4) BR x JP (n=36), 5) BR x JP_we-_ (n=29), 6) BR x JP^rif+^ (n = 13) and 7) BR x JP_we-_^rif+^ (n=5). Each experimental colony was set up with one dark queen pupa, one freshly hatched male, 10 workers and brood (10 eggs, 10 larvae, 10 worker pupae). All individuals were collected from un-manipulated stock colonies, except for JP^rif+^ and JP_we-_^rif+^ males used in combinations #6 and #7, which were collected from rifampicin-treated colonies. To avoid manipulation effects, all combinations were set up in parallel. In such experimental colonies, mating between queens and males usually occurs within three weeks, after which queens shed their wings and being laying eggs. We monitored colonies for queen wing loss once per week. Once queens had shed their wings, we removed any brood placed in the colony during set up and standardized worker number to 20. Colonies were then monitored once a week for six weeks and all eggs and pupae (queen, worker, wingless male, winged male) counted. A subset of colonies was monitored for a total of 12 weeks (JP x BR: n=19, BR x JP: n=28, BR x JP_we-_: n=25, BR x JP^rif+^: n=9, BR x JP_we-_^rif+^: n=5). In each colony, the number of adult workers was kept constant at 20 individuals by removing hatched workers or adding workers from stock colonies. From monitoring data, we calculated mean and maximum weekly egg numbers, and total number and sex ratio of produced pupae (male pupae:total pupae).

To verify that CI is specifically induced by the *w*Cobs-JP strain when it encounters the *w*Cobs-BR strain, we set up three additional mating combinations between queens and males using a third population from Taiwan (TW) which also harbors the *w*Cobs-JP strain: 1) JP x TW (n=5), 2) TW x JP (n=6), 3) BR x TW (n=5). Experimental colonies were set up and monitored as described above for six weeks.

### DNA sequencing and genome assembly

We extracted DNA from a pool of 26 (JP) and 30 (BR) males from two colonies, collected in 2009 in Una, Brazil (BR2009-alpha) from aborted fruits of coconut trees and in 2010 in Naha, Japan (JP201 0-OypB) from under bark of coral trees *(Erythrina* sp.). For details see ^11^. Briefly, the reads were right-tail clipped (minimum quality of 20) and all reads with undefined nucleotides were removed using a combination of FASTX-Toolkit v0.0.14 (http://hannonlab.cshl.edu/fastx_toolkit/, last accessed December 3, 2010) and PRINSEQ++ v1.2 (10.7287/peerj.preprints.27553v1). The resulting paired end reads were assembled with SPAdes v3.13.1 (-k 33,55,77 --only-assembler^66^. The resulting contigs were then binned according to an *ad hoc* BLASTX-based method. First, a proteome database was built from representative genomes (Table S7). Second, the best hit for each scaffold was used for assigning a preliminary bin and a coverage cut-off (10x for BR and 30x for JP) was implemented based on the coverage of the longest contigs. Third, each scaffold was manually inspected using the online NCBI web-server *vs.* BLAST+2.9.0. If the best hits to proteins were consistently to *Wolbachia* bacteria across the scaffold, the query sequence was retained. Finally, the assembly graph was loaded into Bandage v0.8.1^67^, the graph(s) matching *Wolbachia* sequences were identified and the sequences were extracted. These extracted sequences were then compared to the *Wolbachia* draft bin. In both cases, the extracted sequences were roughly the same compounded length as the manually-curated bins and added only a couple of kilo base pairs (kbp), suggesting near completeness of the genome. Using these *Wolbachia* bins, we performed a mapping using BOWTIE2 v2.3.4.1^68^ followed by a re-assembly of the mapped reads in SPAdes (with previously-mentioned parameters). This final reassembly was filtered by both coverage and manual screening of scaffolds shorter than 3 kbp. Assembled sequences have been deposited in the European Nucleotide Archive (ENA) under the accessions CACTIU010000000.1 (*w*Cobs-BR) and CACTIV010000000.1 (*w*Cobs-JP).

### Phylogeny and comparative genomics of *Wolbachia* strains

A draft annotation of the scaffolds was done using Prokka v1.14^69^. For comparative genomic purposes, we built a subset of the predicted ORFs by filtering out all proteins annotated as “hypothetical proteins” with genes shorter than 300 base pairs (bp). This helped us avoid wrongly annotated ORFs overlapping pseudogenes. To infer the phylogenetic relationship between *Wolbachia* strains found to infect the analysed BR and JP populations as well as among other *Wolbachia* bacteria, we collected the amino acid sequences for the ribosomal proteins of 23 *Wolbachia* strains from supergroups A (8), B (10), C (2), D (1), E (1), and F (1) (Table S8). Using *Anap/asma phagocytophilum* strain HZ and *Ehrlichia canis* strain Jake as outgroups, we inferred a Bayesian phylogeny using MrBayes v3.2.6^70^. We ran two independent analyses with four chains each for 300,000 generations and checked for convergence. The substitution model JTT+I + G + F was used as suggested by jModelTest v2.1.10.

To infer the shared protein-coding gene content of *w*Cobs-BR and *w*Cobs-JP strains, we used OrthoMCL v2.0.9^71,72^ with an inflation value of 1.5. The synteny of both strains was assessed using nucmer v3.23^73^ using anchor matches that were unique in both the reference and query (--mum).

### Cif-family gene annotation and phylogenetics

*cif* genes were first identified by BLASTP using the protein sequences WP_010962721.1 (*cifA*) and WP_010962722.1 (*cifB*) as references. Predicted start codons were manually checked for the presence of a Shine-Dalgarno-like sequence. To identify whether the *cif* genes form each strain belonged to the same type or not, we performed a phylogenetic analysis following Lindsey *et. al.* 2018. Briefly, the previously used protein sequences were aligned with the newly acquired ones from both wCobs-BR and wCobs-JP. The alignments were then back-translated to nucleotides. For *cifB*, the type IV genes were excluded and the 3’-end of the alignment was manually truncated to remove the longer unmatched regions of some *cifB* variants. Bayesian phylogenetic inference was performed with MrBayes v3.2.6 with the GTR+G substitution model (as suggested by jModelTest) running two independent analyses with four chains each for 1,000,000 generations, and checked for convergence. Following^23^, Cif protein domains were searched for using HHpred’s v3.2.0 webserver (https://toolkit.tuebingen.mpg.de/tools/hhpred) with default parameters and the following databases: SCOPe70 v2.07, COG/KOG v1.0, Pfam-A v32.0, and SMART v6.0. hits with a probability >=80% were considered.

All genomic data files are available at the zenodo repository https://doi.org/10.5281/zenodo.3561160

### Histological sectioning and fluorescence in situ hybridization

*C. obscurior* (JP) male abdomens were fixed in 4% formaldehyde in PBS and washed with water followed by a dehydration series in n-Butanol (30%, 50%, 70%, 80%, 90%, 96%) at room temperature with shaking for 1 h at each step. Absolute n-butanol was used for the last three hours at 30°C, exchanging the solution each hour. Dehydrated samples were embedded in Technovit 8100 cold polymerizing resin (Heraeus Kulzer, Germany) according to the manufacturer’s instructions, cut into semithin sections (8 μm) with a Microm HM355S microtome (Thermo Fisher Scientific, Germany) and mounted on glass slides coated with poly-L-lysine (Kindler, Germany). The tissue sections were incubated for 90 min at 50°C in hybridization buffer (0.9 M NaCl, 20 mM Tris/HCl (pH 8.0), 0.01 % SDS) containing 0.5 μM of the *Wolbachia*-specific probes Wolb_W2_Cy5 (5’-CTTCTGTGAGTACCGTCATTATC-3’)^74^ and Wolb_Wol3_Cy5 (5’-TCCTCTATCCTCTTTCAATC-3’),^75^, as well as the *Cand*. Westeberhardia-specific probe Wcard1_Cy3 (5’-ATCAGTTTCGAACGCCATTC-3’)^10^. DAPI (4’,6-diamidino-2-phenylindole) was used for host DNA counterstaining. After hybridization, samples were washed with buffer (0.1 M NaCl, 20 mM Tris/HCl (pH 8.0), 5mM EDTA, 0.01 % SDS) for 20 min at 50°C. Subsequently, the washing buffer was removed and washed with distilled water for 20 min at 50°C. Samples were air-dried and covered with VectaShield^®^ (Vector Laboratories, Burlingame, CA, USA) and stored overnight at room temperature. Images were acquired on an AxioImager.Z1 epifluorescence microscope (Carl Zeiss, Jena, Germany).

## Contributions

CÜ performed the crosses and qPCR

ES analyzed and visualized the data

AMM assembled *Wolbachia* genomes, performed comparative genomic analyses, calculated the phylogeny and annotated CI genes

LVF conducted FISH analyses

BS revised the taxonomic status of the populations

JH acquired funding and reviewed and edited the manuscript

JO conceived and supervised the study

JO, ES and CÜ wrote the original draft of the manuscript

All authors revised and approved the final version of the manuscript

## Acknowledgements

The authors thank J. Wang for help collecting colonies in Taiwan and for providing the *wsp* sequence of the TW population. We thank J. Hacker for help with the BR x TW crossings and H. Lowack and J. Wallner for help with lab work. J.D. Shropshire helped with the *cif*annotation and provided useful comments on a previous version of the manuscript. B. Weiss assisted with histological preparations. This work was supported by the Marie-Curie AgreenSkills+ fellowship program cofunded by the EU’s Seventh Framework Programme (FP7-609398) to A.M.M, and a DFG grant He1 623/31 to J.H. and J.O.

