## Supplement Figures and Tables for "Cytoplasmic incompatibility between Old and New World populations of a tramp ant"

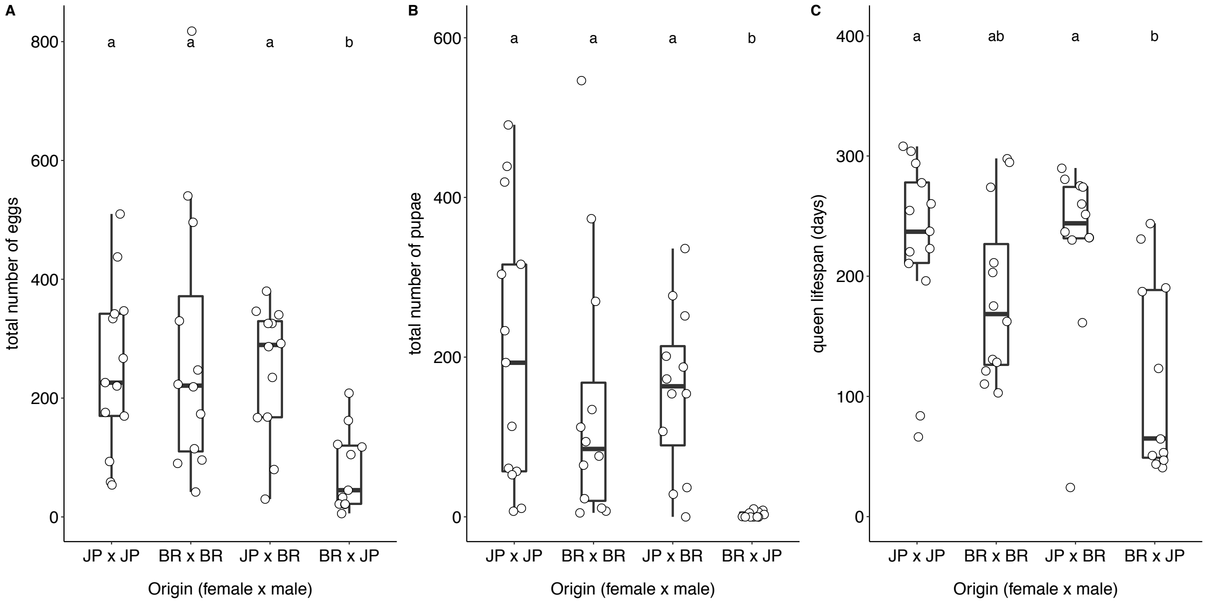


Figure S1: Cytoplasmic incompatibility between populations consistently reduces colony fitness

Cytoplasmic incompatibility between Brazilian (BR) and Japanese (JP) ants consistently reduces lifetime colony productivity monitored twice per week (A, B) and queen lifespan (C). BR queens mated to JP males (BR x JP) produced fewer eggs (A) and pupae (B) than the reciprocal cross JP x BR and the intrapopulation crosses JP x JP and BR x BR. BR queens mated to JP males also exhibit shortened lifespans (C) compared to JP x BR and JP x JP crosses. Experimental crosses were set up from independent stock colonies with one queen pupa, one male pupa and 20 workers (JPxJP: n=13, BRxBR: n=12, JPxBR: n=12, BRxJP: n=11). Differences between groups were tested with pairwise Mann-Whitney-U-tests followed by Bonferroni-Holms correction of p-values. Letters above boxplots indicate statistically significant differences at $p<0.05$.

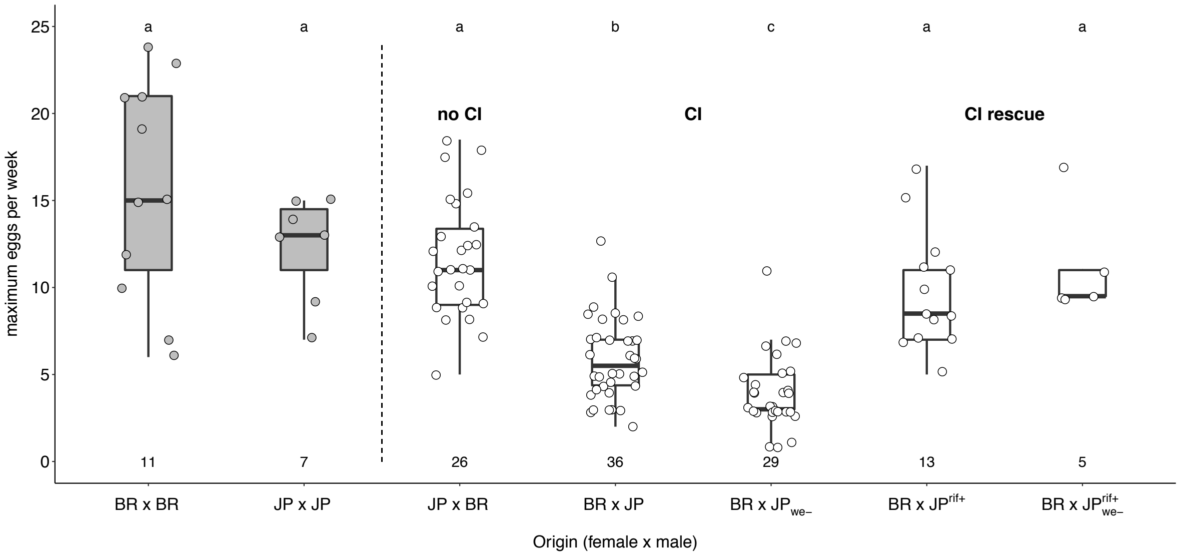


Figure S2: Maximum weekly egg numbers produced by inter-population crosses Maximum weekly egg numbers produced by Brazilian (BR) and Japanese (JP) queens mated to males from their own or from a different population. JP ants were collected from colonies that either carried the main endosymbiont *Candidatus* Westeberhardia cardiocondylae (JP) or did not (JP$_{we-}$). Males used in "CI rescue" crosses were collected from JP colonies with and without *Cand.* Westeberhardia cardiocondylae that had been treated with the antibiotic rifampicin (JP$^{rif+}$; JP$_{we-}^{rif+}$). Numbers below box plots indicate the number of replicates. Differences between groups were tested with pairwise Mann-Whitney-U-tests followed by Bonferroni-Holms correction of p-values. Letters above boxplots indicate statistically significant differences at $p<0.05$. For Bonferroni-Holms corrected pairwise p-values see Table S2.

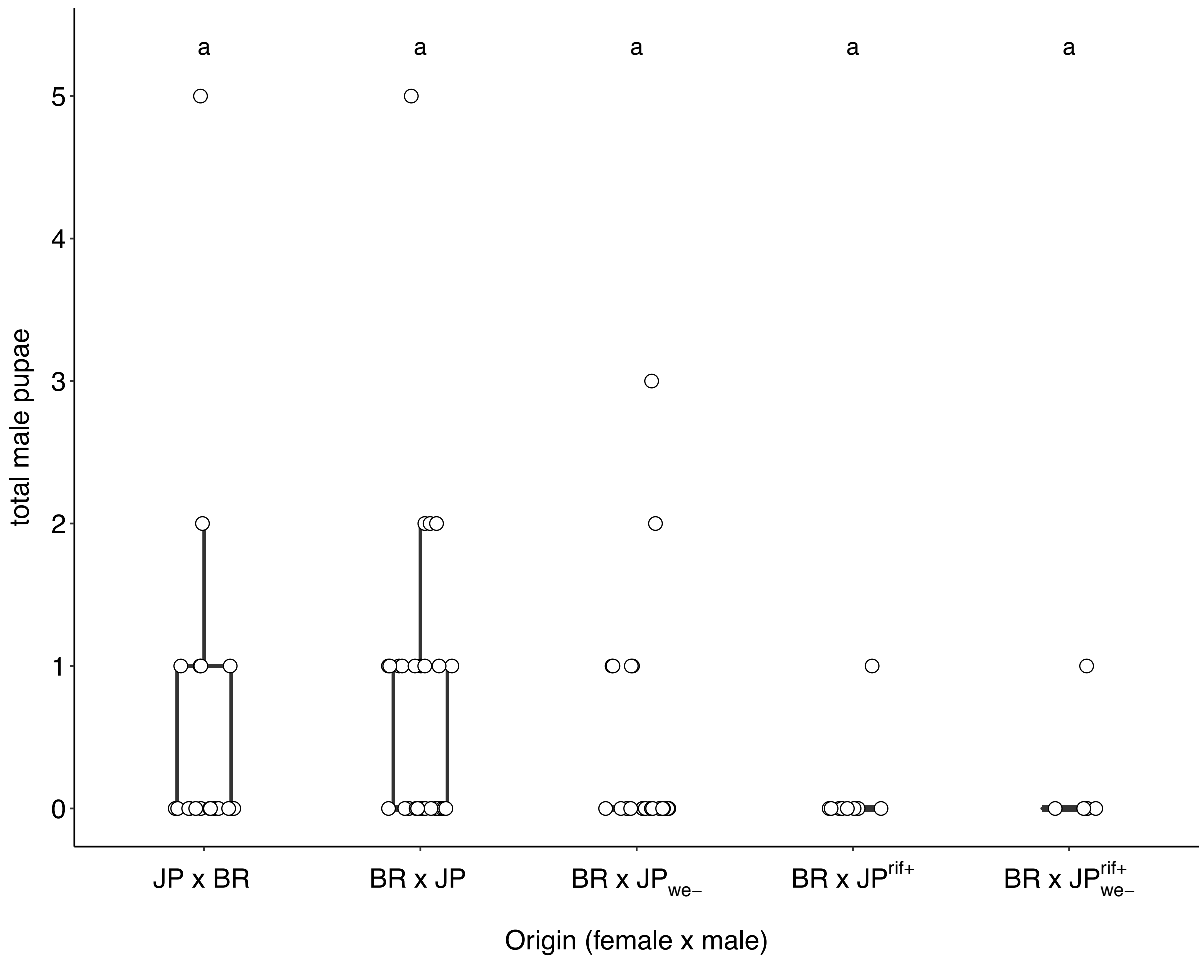


Figure S3: Total number of male pupae produced in inter-population crosses
Total number of male pupae produced in crosses between Brazilian (BR) and Japanese (JP) males and queens monitored for 12 weeks. JP ants were collected from colonies that either carried the main endosymbiont *Candidatus* Westeberhardia cardiocondylae (JP) or did not (JP$_{we-}$). Males used in "CI rescue" crosses were collected from JP colonies with and without *Cand.* Westeberhardia cardiocondylae, which had been treated with the antibiotic rifampicin (JP$^{rif+}$; JP$_{we-}^{rif+}$). Differences between groups were tested with pairwise Mann-Whitney-U-tests followed by Bonferroni-Holms correction of p-values. Letters above boxplots indicate statistically significant differences at $p<0.05$.

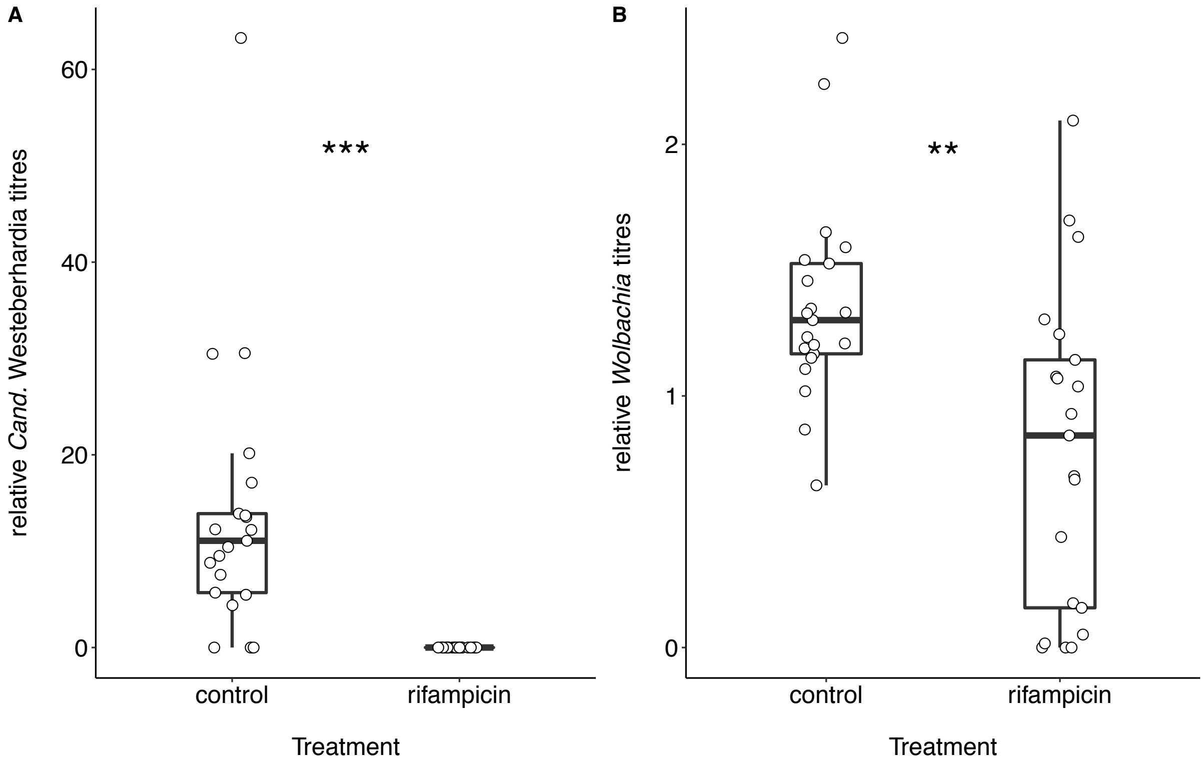


Figure S4: Effects of rifampicin treatment on endosymbiont titres in worker pupae
Rifampicin treatment significantly decreases *Cand*. Westeberhardia cardiocondylae (A) and *Wolbachia* (B) titres in worker pupae. All pupae were collected from JP colonies that carried the main endosymbiont *Candidatus* Westeberhardia cardiocondylae. Differences in endosymbiont titres between control and rifampicin-treated ants were tested with Mann-Whitney-U-Tests. Stars indicate statistically significant differences at $p<0.01$ (**) and $p<0.001$ (***).

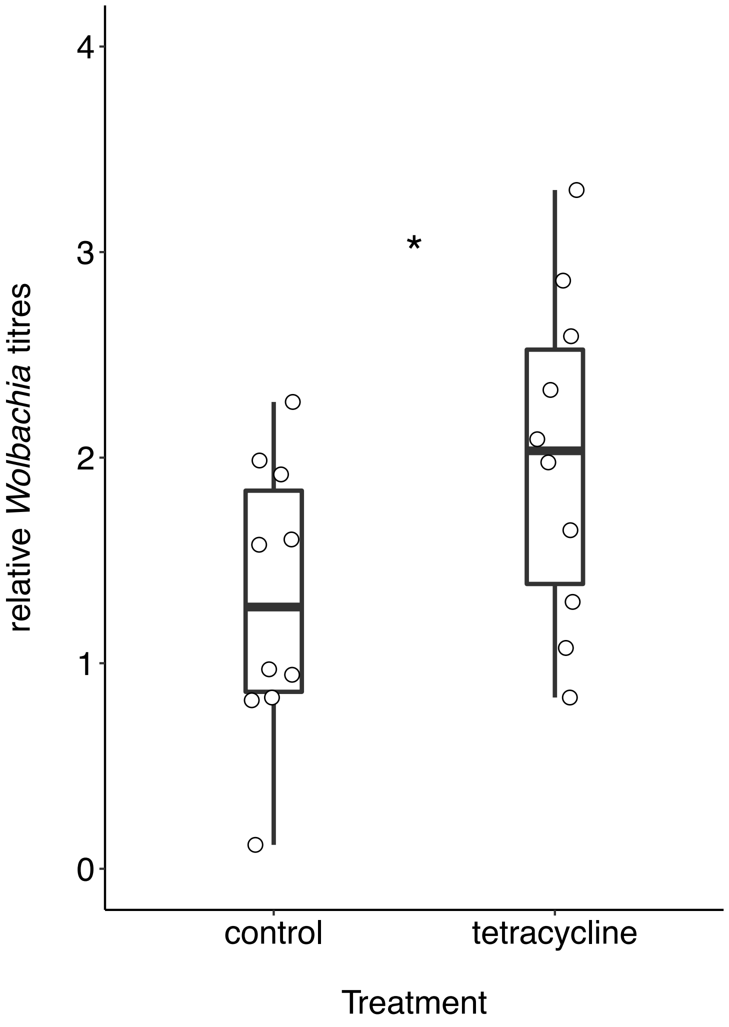


Figure S5: Effects of tetracycline treatment on *Wolbachia* titres in workers
Tetracycline treatment (0.5% in a 1:1 honey-water solution) increases *Wolbachia* titres in worker pupae. Tetracycline treatment followed the same procedure as rifampicin treatment. Differences in endosymbiont titres between control and tetracycline-treated ants were tested with a Mann-Whitney-U-Test. Stars indicate statistically significant differences at $p<0.05$ (*).

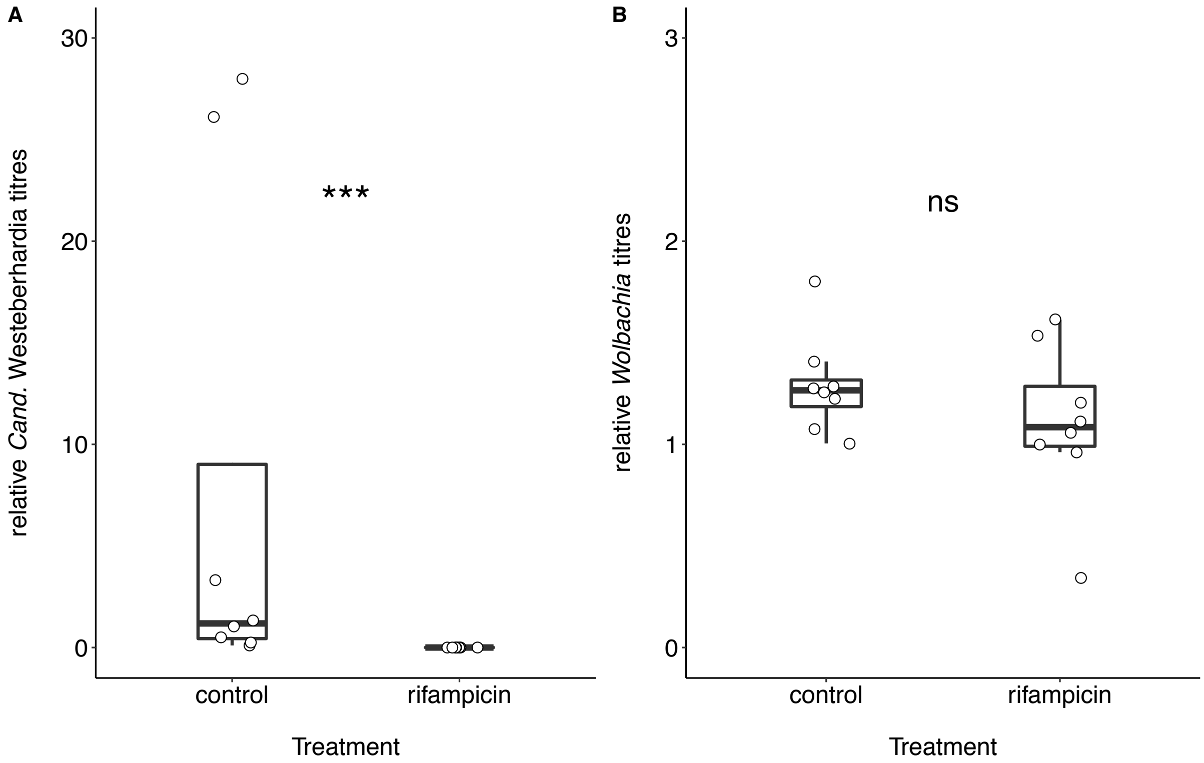


Figure S6: Recovery of endosymbiont titres after rifampicin treatment
Recovery of *Cand.* Westeberhardia cardiocondylae (A) and *Wolbachia* (B) titres in workers six months after rifampicin treatment. All workers were collected from JP colonies that previously carried the main endosymbiont *Candidatus* Westeberhardia cardiocondylae. Differences in endosymbiont titres between control and rifampicin-treated ants were tested with Mann-Whitney-U-Tests. Stars indicate statistically significant differences at p<0.001 (***).

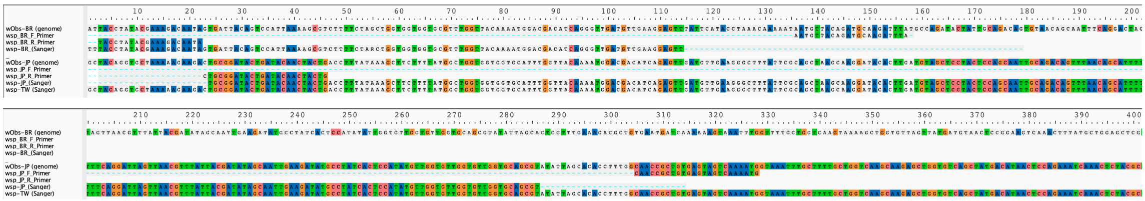

Figure S7: *Wolbachia surface protein* sequences differ between populations
Alignment of *Wolbachia surface protein* (*wsp*) primers and sequences obtained from *C. obscurior* samples collected in Brazil (BR), Japan (JP) and Taiwan (TW). Sequences were extracted from whole genomes (BR: sequence #1, JP: sequence #5) and Sanger-sequenced PCR products (BR: sequence #4, JP: sequence #8, TW: sequence #9). BR and JP sequences differ in 21 nucleotides while the sequence obtained from TW samples shows 100% sequence identity with JP sequences.

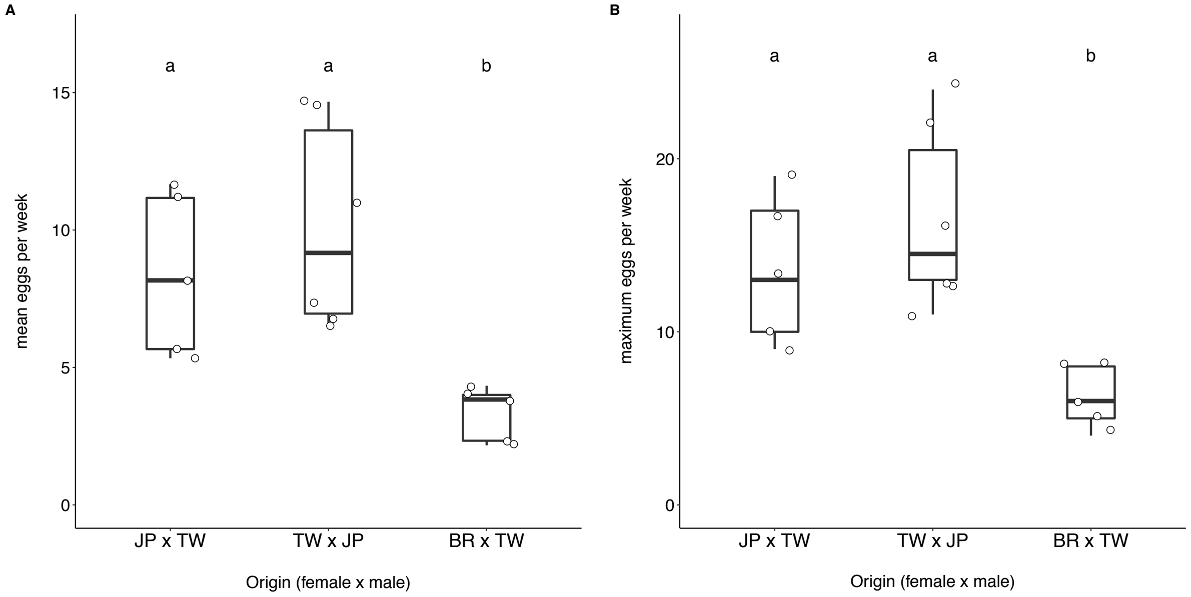


Figure S8: Unidirectional cytoplasmic incompatibility between populations
Mean (A) and maximum (B) weekly egg numbers produced by crosses between queens and males from Taiwan (TW), Japan (JP) and Brazil (BR). Differences between groups were tested with Kruskal Wallis rank sum tests followed by pairwise Mann-Whitney-U-tests and Bonferroni-Holms correction of p-values. Letters above boxplots indicate statistically significant differences at $p<0.05$.

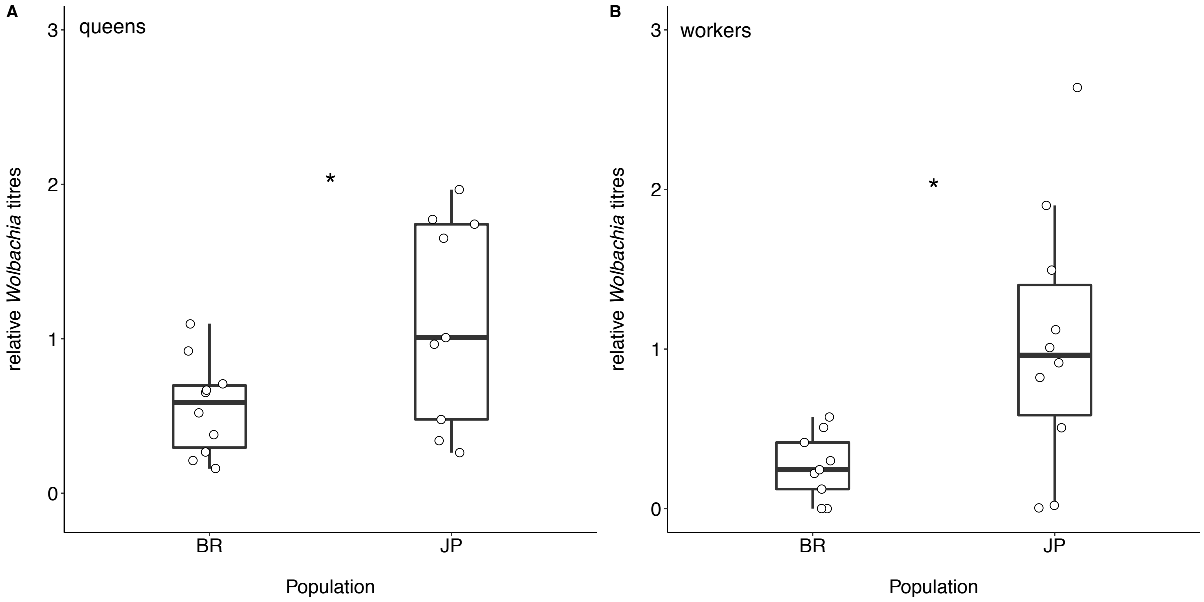


Figure S9: Population-specific *Wolbachia* titres
*Wolbachia* titres in queens (A) and workers (B) are significantly lower in colonies collected in Brazil (BR) compared to colonies collected in Japan (JP). Differences in *Wolbachia* titres between populations were tested with Mann-Whitney-U-Tests. Stars indicate statistically significant differences at $p<0.05$ (*).

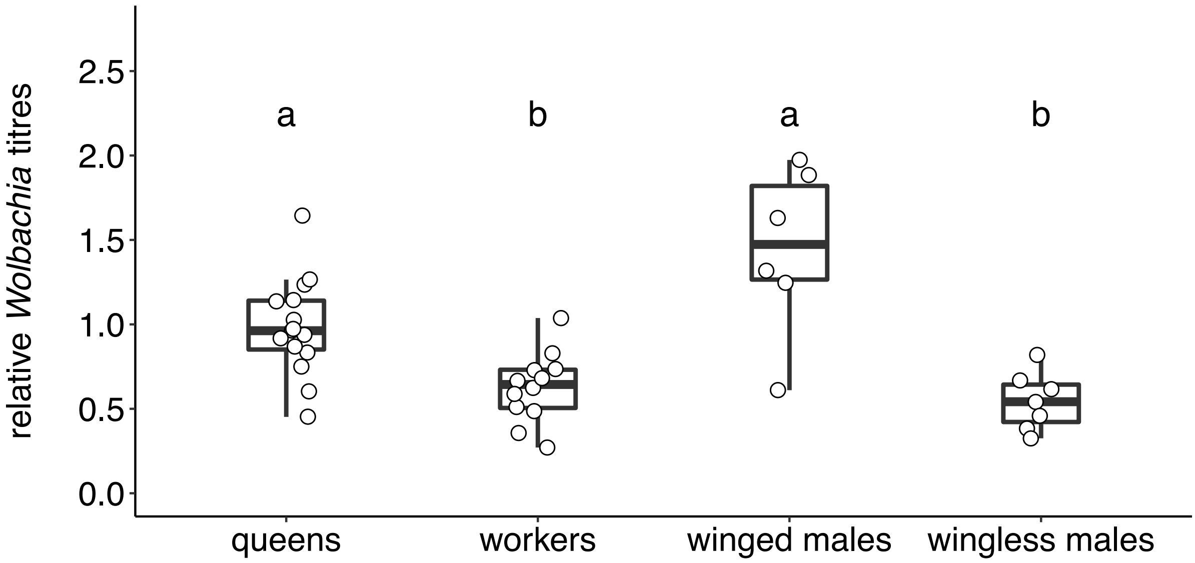


Figure S10: Morph-specific *Wolbachia* titres
Differences between groups were tested with a Kruskal Wallis rank sum test followed by pairwise Mann-Whitney-U-tests and Bonferroni-Holms correction of p-values. Letters above boxplots indicate statistically significant differences at $p<0.05$.

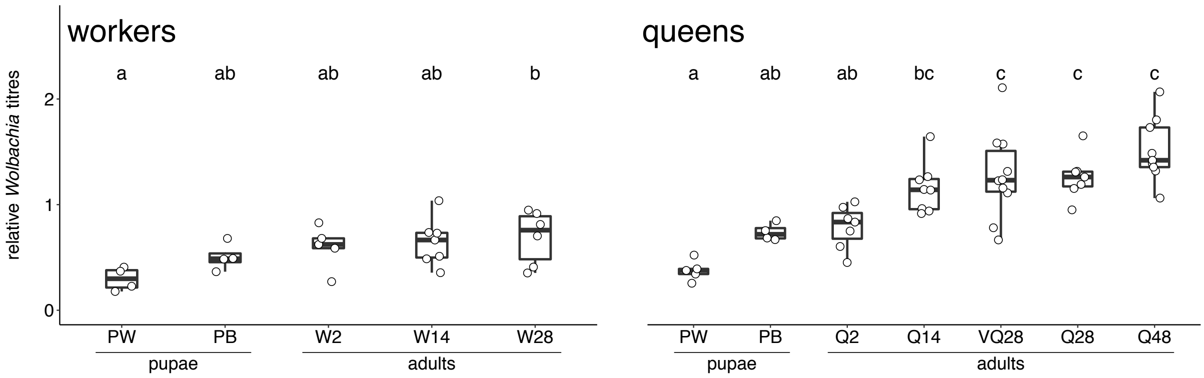


Figure S11: Age-dependent changes in *Wolbachia* titres in queens and workers *Wolbachia* titres were measured in pupae (PW: white pupa, PB: brown pupa) and adults of different ages (W2/Q2: two day-old worker/queen; W14/Q14: 14 day-old worker/queen; W28/VQ28/Q28: 28 day-old worker/virgin queen/queen; Q48: 48 day-old queen). For each morph (workers, queens), differences between development stages and ages were tested with ANOVAs followed by Tukey post-hoc tests for p-value correction. Letters above boxplots indicate statistically significant differences at $p<0.05$.

Supplementary Tables

Table S1: Bonferroni-Holms corrected p-values for pairwise comparisons of mean egg numbers produced by crosses between queens and males from Brazil (BR) and Japan (JP) during the first 6 weeks

| | BR x BR | JP x JP | JP x BR | BR x JP | BR x $JP_{we-}$ | BR x $JP^{rif+}$ | BR x $JP_{we-}^{rif+}$ |
|---|---|---|---|---|---|---|---|
| BR x BR | - | | | | | | |
| JP x JP | 1 | - | | | | | |
| JP x BR | 0.7454 | 1 | - | | | | |
| BR x JP | 0.0013 | 0.0032 | 4.3e-06 | - | | | |
| BR x $JP_{we-}$ | 6.2e-05 | 0.0009 | 7.1e-08 | 0.0086 | - | | |
| BR x $JP^{rif+}$ | 0.6363 | 0.7891 | 1 | 0.0024 | 1.6e-05 | - | |
| BR x $JP_{we-}^{rif+}$ | 1 | 1 | 1 | 0.0167 | 0.0059 | 0.6363 | - |

Table S2: Bonferroni-Holms corrected p-values for pairwise comparisons of maximum egg numbers produced by crosses between queens and males from Brazil (BR) and Japan (JP) during the first 6 weeks

| | BR x BR | JP x JP | JP x BR | BR x JP | BR x $JP_{we-}$ | BR x $JP^{rif+}$ | BR x $JP_{we-}^{rif+}$ |
|---|---|---|---|---|---|---|---|
| BR x BR | - | | | | | | |
| JP x JP | 1 | - | | | | | |
| JP x BR | 0.6024 | 1 | - | | | | |
| BR x JP | 0.0004 | 0.0032 | 1 | - | | | |
| BR x $JP_{we-}$ | 6.5e-05 | 0.0012 | 2.8e-08 | 0.0055 | - | | |
| BR x $JP^{rif+}$ | 0.2914 | 0.8393 | 0.6024 | 0.0043 | 5.4e-05 | - | |
| BR x $JP_{we-}^{rif+}$ | 1 | 1 | 1 | 0.0105 | 0.0086 | 1 | - |

Table S3: Bonferroni-Holms corrected p-values for pairwise comparisons of total pupae numbers produced by crosses between queens and males from Brazil (BR) and Japan (JP) during 12 weeks

| | JP x BR | BR x JP | BR x $JP_{we-}$ | BR x $JP^{rif+}$ | BR x $JP_{we-}^{rif+}$ |
|---|---|---|---|---|---|
| JP x BR | - | | | | |
| BR x JP | 4.1e-06 | - | | | |
| BR x $JP_{we-}$ | 3.7e-06 | 1 | - | | |
| BR x $JP^{rif+}$ | 0.091 | 0.081 | 0.091 | - | |
| BR x $JP_{we-}^{rif+}$ | 1 | 0.047 | 0.064 | 1 | - |

Table S4: Bonferroni-Holms corrected p-values for pairwise comparisons of sex ratios produced by crosses between queens and males from Brazil (BR) and Japan (JP) during 12 weeks

| | JP x BR | BR x JP | BR x $JP_{we-}$ | BR x $JP^{rif+}$ | BR x $JP_{we-}^{rif+}$ |
|---|---|---|---|---|---|
| JP x BR | - | | | | |
| BR x JP | 3.490e-06 | - | | | |
| BR x $JP_{we-}$ | 0.0577 | 0.9395 | - | | |
| BR x $JP^{rif+}$ | 1 | 0.0368 | 0.3164 | - | |
| BR x $JP_{we-}^{rif+}$ | 1 | 0.0839 | 0.5884 | 1 | - |

Table S5: Bonferroni-Holms corrected p-values for pairwise comparisons of mean egg numbers produced by crosses between queens and males from Brazil (BR), Japan (JP), and Taiwan (TW) during the first 6 weeks

| | JP x TW | TW x JP | BR x TW |
|---|---|---|---|
| JP x TW | - | | |
| TW x JP | 0.537 | - | |
| BR x TW | 0.016 | 0.013 | - |

Table S6: Bonferroni-Holms corrected p-values for pairwise comparisons of maximum egg numbers produced by crosses between queens and males from Brazil (BR), Japan (JP), and Taiwan (TW) during the first 6 weeks

| | JP x TW | TW x JP | BR x TW |
|---|---|---|---|
| JP x TW | - | | |
| TW x JP | 0.407 | - | |
| BR x TW | 0.024 | 0.023 | - |

Table S7: Organisms and accession numbers used for BLASTX binning of assembled scaffolds

| Species | Strain / isolate | Accession / link | Target organism |
|---|---|---|---|
| Cardiocondyla obscurior | - | https://www.antgenomes.org/ | Ant host |
| Cardiocondyla obscurior | COBS20161003 | KX951753.1 | Ant mitochondrion |
| Ca. Westeberhardia cardiocondylae | obscurior BR-alpha | LN774881.1 | Cand. Westeberhardia cardiocondylae |
| Wolbachia sp. | wMel | AE017196.1 | Wolbachia |
| Wolbachia sp. | TRS | AE017321.1 | Wolbachia |
| Wolbachia sp. | wPip | AM999887.1 | Wolbachia |
| Wolbachia sp. | wCle | AP013028.1 | Wolbachia |
| Wolbachia sp. | wRi | CP001391.1 | Wolbachia |
| Wolbachia sp. | wNo | CP003883.1 | Wolbachia |
| Wolbachia sp. | wInc_Cu | CP011148.1 | Wolbachia |
| Wolbachia sp. | Berlin | CP015510.2 | Wolbachia |
| Wolbachia sp. | China 1 | CP016430.1 | Wolbachia |
| Wolbachia sp. | wAlbB | CP031221.1 | Wolbachia |
| Wolbachia sp. | WOo | HE660029.1 | Wolbachia |
| Wolbachia sp. | wTpre | LKEQ01000000.1 | Wolbachia |
| Wolbachia sp. | Cameroon | NZ_HG810405.1 | Wolbachia |

Table S8: Organism details of *Wolbachia* strains and outgroup bacteria used for Bayesian phylogenetic placement of newly sequenced *Wolbachia* strains.

| Species | Strain / isolate / voucher | Accession / link | Host |
|---|---|---|---|
| *Anaplasma phagocytophilum* | HZ | CP000235.1 | Homo sapiens |
| *Ehrlichia canis* | Jake | CP000107.1 | *Canis lupus familiaris* |
| *Wolbachia* sp. | Berlin | CP015510.2 | *Folsomia candida* |
| *Wolbachia* sp. | TRS | AE017321.1 | *Brugia malayi* |
| *Wolbachia* sp. | wCle | AP013028.1 | *Cimex lectularius* |
| *Wolbachia* sp. | WOo | HE660029.1 | *Onchocerca ochengi* |
| *Wolbachia* sp. | Cameroon | NZ_HG810405.1 | *Onchocerca volvulus* |
| *Wolbachia* sp. | wCauA | CP041215.1 | *Carposina sasakii* |
| *Wolbachia* sp. | wHa | CP003884.1 | *Drosophila simulans* |
| *Wolbachia* sp. | wDana W2.1 | CP042904.1 | *Drosophila ananassae* |
| *Wolbachia* sp. | wRi | CP001391.1 | *Drosophila simulans* |
| *Wolbachia* sp. | wAu | LK055284.1 | *Drosophila simulans* |
| *Wolbachia* sp. | wMel | AE017196.1 | *Drosophila melanogaster* |
| *Wolbachia* sp. | wStri | NZ_MUIX01000000.1 | *Laodelphax striatellus* |
| *Wolbachia* sp. | China 1 | CP016430.1 | *Bemisia tabaci* |
| *Wolbachia* sp. | wLug | NZ_MUIY01000000.1 | *Nilaparvata lugens* |
| *Wolbachia* sp. | wAlbB | CP031221.1 | *Aedes albopictus* |
| *Wolbachia* sp. | wTpre | LKEQ01000000.1 | *Trichogramma pretiosum* |
| *Wolbachia* sp. | wMau | CP034335.1 | *Drosophila mauritiana* |
| *Wolbachia* sp. | wNo | CP003883.1 | *Drosophila simulans* |
| *Wolbachia* sp. | GBW | QJHA01000000.1 | *Leptopilina clavipes* |
| *Wolbachia* sp. | wMeg | CP021120.1 | *Chrysomya megacephala* |
| *Wolbachia* sp. | wPip | AM999887.1 | *Culex quinquefasciatus* |
